# Budding yeast Wpl1p regulates cohesin functions in cohesion, condensation and DNA repair by a common mechanism

**DOI:** 10.1101/169474

**Authors:** Michelle S. Bloom, Vincent Guacci, Douglas Koshland

## Abstract

Cohesin tethers DNA to mediate sister chromatid cohesion, chromosome condensation, and DNA repair. How the cell regulates cohesin to perform these distinct functions remains to be elucidated. One cohesin regulator, Wpl1p, was characterized in the budding yeast, *Saccharomyces cerevisiae*, as a promoter of cohesion and as an inhibitor of condensation. Here we provide evidence that Wpl1p has an additional function in promoting the timely repair of DNA damage induced during S-phase. In addition to these biological functions, Wpl1p has been implicated as an inhibitor of cohesin’s ability to stably bind DNA by modulating the interface between two subunits (Mcd1p and Smc3p) of the core cohesin complex. We show that Wpl1p likely modulates this interface to regulate all cohesin’s biological functions. Furthermore, we show that Wpl1p regulates cohesion and condensation through the formation of a functional complex with another cohesin-associated factor, Pds5p. In contrast, Wpl1p regulates DNA repair independently of its interaction with Pds5p. Together these results suggest that Wpl1p regulates distinct biological functions of cohesin by Pds5p-dependent and – independent modulation of the Smc3p-Mcd1p interface.

## Introduction

Cohesin, a member of the SMC family of protein complexes, is comprised of four-subunits, Smc1p, Smc3p, Mcd1p (Scc1/Rad21), and Scc3p (SA/STAG). Cohesin mediates a myriad of nuclear functions essential for both viability and accurate transmission of genetic information including sister chromatid cohesion, condensation of chromosomes, and the repair of DNA damage during G2/M (Onn et al. 2008). Cohesin is thought to perform these different functions through the spatial and temporal regulation of its ability to tether two genomic loci (Guacci et al. 1997; Gruber et al. 2003). Cohesin’s DNA binding and tethering activities are regulated by factors including Eco1p (Ctf7p), Pds5p, and Wpl1p (Rad61p) (Skibbens et al. 1999; Hartman et al. 2000; Rolef Ben-Shahar et al. 2008; Unal et al. 2008). How these regulatory factors interface with each other and with cohesin to promote its biological functions remains poorly understood.

The factor Wpl1p was first implicated as a negative regulator of the cohesin complex, serving to inhibit both cohesion and condensation. Evidence that Wpl1p is an inhibitor of condensation stems from findings that the deletion of *WPL1* (*wpl1*Δ) restores viability and condensation to cells lacking Eco1p function (*eco1*Δ) (Guacci and Koshland 2012; Lopez-Serra et al. 2013) and that *wpl1*Δ in a wild-type background leads to premature condensation as compared to wild-type cells (Lopez-Serra et al. 2013). Additionally, Wpl1p’s role as an inhibitor of cohesion was supported by the loss of cohesion seen when Wpl1p was overexpressed in both human and yeast cells (Gandhi et al. 2006; Lopez-Serra et al. 2013). Wpl1p is thought to inhibit cohesin function by removing it from DNA in a non-proteolytic manner. Recent biochemical studies suggest that Wpl1p destabilizes the interface between the N-terminus of Mcd1p and the base of the coiled-coil of Smc3p (Buheitel and Stemmann 2013; Beckouët et al. 2016). Additionally, mutating an Smc3p residue in the Smc3p/Mcd1p interface abolishes cohesin localization to centromere proximal regions, providing *in vivo* support for a role for this interface (Gligoris et al. 2014). However, the biological function and regulation of destabilization of the Smc3p/Mcd1p interface is poorly understood.

To limit Wpl1p inhibition, cohesin is acetylated by Eco1p at two conserved lysine residues on Smc3p (K112 K113 in the budding yeast, *Saccharomyces cerevisiae*) (Rolef Ben-Shahar et al. 2008; Unal et al. 2008). Additionally, Pds5p helps preserve Smc3p acetylation during and after S-phase, suggesting a common molecular mechanism for how Pds5p and Eco1p promote cohesion (Chan et al. 2013). These functions are thought to promote condensation as inactivation of either factor results in dramatic defects in both processes (Skibbens et al. 1999; Hartman et al. 2000). Further corroboration that Pds5p and Eco1p promote cohesin function through a common molecular mechanism is that over-expression of Pds5p suppresses mutants containing *eco1*-*ts* alleles, and vice-versa (Noble et al. 2006). Taken together, these data suggest that Eco1p and Pds5p both prevent Wpl1p-mediated antagonization of cohesion and condensation.

However, the function of Wpl1p and Pds5p in regulating cohesin is more complicated. In budding yeast, *wpl1*Δ cells surprisingly display a partial cohesion defect, implicating Wpl1p as a positive factor required for efficient cohesion, though the molecular differences between Wpl1p’s positive and negative functions remains a mystery (Rowland et al. 2009; Sutani et al. 2009; Guacci and Koshland 2012). Furthermore, Wpl1p and Pds5p have been shown to form a complex that is capable of unloading of cohesin from DNA *in vitro* (Kueng et al. 2006; Murayama and Uhlmann 2015). This result suggests that Pds5p may inhibit cohesin in addition to its well-established role in promoting cohesin function. Consistent with this idea, in *S. pombe*, the deletion of *pds5* suppresses a deletion of the *ECO1* homologue, Eso1 (Tanaka et al. 2001). Moreover, in budding yeast, certain *PDS5* alleles suppress the inviability of *eco1*-*ts* mutants which have reduced cohesin acetylation (Rowland et al. 2009; Sutani et al. 2009). This suppression is consistent with the idea that these *pds5* mutations inactivate a Pds5p-mediated inhibitory activity. Together these results suggest that Wpl1p and Pds5p can act both positively and negatively to regulate cohesin functions.

The complex regulation of Wpl1p on cohesin function raises important questions that we address in this study. First, are there additional roles of Wpl1p in regulating cohesin function? Does Wpl1p regulate all cohesin’s biological functions through a common molecular mechanism? Finally, Is Wpl1p’s ability to form a complex with Pds5p important for any or all of Wpl1p’s regulatory functions? The answers to these questions provide important new insights into cohesin regulation by Wpl1p and its interplay with Pds5p.

## Materials and Methods

### Yeast strains, media, and reagents

Yeast strains used in this study are A364A background, and their genotypes are listed in Table 2.1. YPD liquid media was prepared containing 1% yeast extract, 2% peptone, 2% dextrose, 0.01 mg/ml adenine.

#### Solid Media

YPD solid media was prepared containing 1% yeast extract, 2% peptone, 2% dextrose, 2% agar.

#### Camptothecin

Camptothecin (Sigma-Aldrich, St. Louis, MO) was made as a 10 mg/ml stock (in DMSO) and added to final concentration of 20 μg/ ml in YPD media containing 25 mM pH 7.4 4-(2-hydroxyethyl)-1-pipera-zineethanesulfonic acid (HEPES; Fisher Scientific, Fair Lawn, NJ).

#### Methyl-methane sulfonate

99% pure MMS (Sigma-Aldrich, St. Louis, MO) was added to a final concentration of 0.01% in YPD media. MMS was diluted 1:10 in dimethyl sulfoxide (DMSO) for addition to liquid media. Agar plates containing MMS were made within 2 days of use to prevent degradation.

#### Dropout media

5-FOA was purchased from US Biological Life Sciences (Salem, MA) and used at a final concentration of 1 g/L in URA dropout plates supplemented with 50 mg/L uracil powder (Sigma-Aldrich)

### Dilution plating

Cells were grown to saturation in YPD liquid media at 30°C then plated in 10-fold serial dilutions. Cells containing temperature-sensitive alleles were grown to saturation at 23°C. Cells were incubated on plates at relevant temperatures or containing drugs as described. For plasmid shuffle assays, cells were grown to saturation in YPD media to allow loss of covering plasmid, then plated in 10-fold serial dilutions on YPD or FOA media.

#### Cohesin and condensation time course

Cells are inoculated into 5 mL YPD starter culture overnight at 23°C, unless indicated otherwise. Cells are then inoculated from starter cultures into YPD to grow overnight to a final concentration of 0.2 OD. Alpha factor (Sigma-Aldrich) is added to cultures at 10^−8^ M, for 3 hours for cells to arrest in G1. Cells are then washed 3x in YPD containing .2 μg/mL Pronase E, and washed 1x in YPD without Pronase E. Cells are then resuspended into YPD containing 15 μg/mL nocodazole (Sigma-Aldrich) and incubated at 23°C to allow cell cycle progression until arrest in mid-M (3 hours). To assess cohesion through separation of LacI-GFP foci, cells were fixed for 15-30 minutes in 4% paraformaldehyde (w/v) 3.4% sucrose (w/v) solution, and then washed and resuspended in 0.1 KPO_4_ 1.2 M sorbitol buffer then stored at 4°C.

For auxin treatment, time courses were performed as above except 1M 3-indoleacetic acid (Sigma-Aldrich, St. Louis, MO) was dissolved in dimethyl sulfoxide (DMSO) and added to final concentration of 500 μM to alpha factor arrested cells, then incubated an additional 1 hour. 500 μM auxin was present in all YPD washes and the releasing media containing nocodazole.

### CPT and MMS treatment time course

Cells were grown, arrested in G1, and released as described above. Upon release from alpha-factor, cells were split and resuspended into YPD containing either DMSO, 20 μg/mL CPT and 25 mM HEPES pH7.4 or 0.01% MMS and incubated at 23°C to allow cell cycle progression. 90 minutes after release, alpha-factor was re-added to cultures at 10^−8^ M to arrest in subsequent G1. Cells were harvested every 30-minutes and fixed in 70% ethanol. To assess chromosome segregation, fixed cells were washed and resuspended in 1×PBS containing DAPI.

Assessment of chromosome segregation when treated with MMS or CPT in nocodazole (G2/M) cells. Cells were grown, arrested and released from alpha factor arrest into nocodazole in the absence of drugs as described above. Cells were then released from nocodazole arrest by washing 3x in YPD. Cells were then split and resuspended into media containing alpha-factor at 10^−8^ M and either DMSO, 20 μg/mL CPT and 25 mM HEPES pH 7.4 or 0.01% MMS.

### Fluorescence in situ hybridization

Nocodazole arrested cells were fixed in 3.6% formaldehyde for 2 hours at 23°C. Cells were then washed 3x with water and resuspended in 1M sorbitol 20mM KPO_4_ pH 4.7. Cells were spheroplasted with beta mercaptoethanol and 0.5% TritonX-100. Cells were then gently spun down and plated on polylysine-coated slides. Cells were then washed with 0.5% SDS. Slides were then submerged in 3:1 methanol/acetic acid and allowed to air-dry overnight. Cells were then RNase A treated (100 μg/mL in 2X SSC) at 37% for 1 hr. Slides were washed 4x in fresh 2X SSC and immediately dehydrated through a series of ethanol washes (70%, 80% and 95%, min/wash at -20°C). Denaturation of chromosomal DNA was done by incubation of slides in 70% formamide in 2X SSC at 70°C for 2 minutes, followed immediately by ethanol washes as described above (70%, 80%, 90%, and 100%). After slides had dried, cells were treated with proteinase K (10 μg/mL in 20 mM Tris, pH 7.2, 2 mM CaCl2) for 15 minutes at 37°C. Cells were then stained with ProLong Gold Antifade Mountant with DAPI (Life Technologies).

#### Flow cytometry

To assess DNA content, cells were fixed in 70% ethanol. Fixed cells were washed twice in 50 mM sodium citrate (pH 7.2), then treated with RNase A (50 mM sodium citrate [pH 7.2]; 0.25 mg/ml RNase A; 1% Tween-20 [v/v]) overnight at 37°C. Proteinase K was then added to a final concentration of 0.2 mg/ml and samples were incubated at 50°C for 2 hr. Samples were sonicated for 30s or until cells were adequately disaggregated. SYBR Green DNA I dye (Life Technologies, Carlsbad, CA) was then added at 1:20,000 dilution and samples were run on a Guava easyCyte flow cytometer (Millipore, Billerica, MA). 20,000 events were captured for each time point. Quantification was performed using FlowJo analysis software.

### Microscopy

Images were acquired with an Axioplan2 microscope (100x objective, numerical aperture [NA] 1.40; Zeiss, Thornwood, NY) equipped with a Quantix charge-coupled device camera (Photometrics, Tucson, AZ).

### Preparation of cells for immunoprecipitation

Cells were inoculated into 5 mL starter cultures and grown overnight at 23°C. Strains were then inoculated into 60 mL cultures and grown to a final OD of 0.8. 20 ODs were then harvested, washed in 1XPBS, spun down, liquid aspirated and cells were flash frozen in LN_2_.

For CPT, untreated cells are grown to a final OD_600_ of 0.4. 1M HEPES pH 7.4 is added to cultures to a final concentration of 25 mM. 10 mg/mL CPT stock is added to cells to final concentration of 20 μg/ml and cells are incubated for 3 hours. 20 OD were then harvested and prepared as described above.

### Immunoprecipitation

Cell lysates were prepared by bead beating 30 sec on 1 min rest, 4x at 4°C in GNK100 buffer (100 mM KCl, 20 mM HEPES pH 7.5, 0.2% NP40, 10% glycerol, 2.5 mM MgCl_2_) containing complete mini EDTA free protease inhibitor (Roche), 5mM sodium butyrate, 5 mM beta mercaptoethanol, 1 mM PMSF, and 20 mM b-glycerophosphate. Lysates were cleared of insoluble cell debris and then incubated with anti-FLAG antibody (Sigma-Aldrich, St. Louis, MO) and Protein A dynabeads for 1 hour at 4°C. Dynabeads were then washed 4x with GNK100 buffer with additives as described above containing 100 μM MG132. Samples were then run on SDS-PAGE gels and analyzed through western blot.

## Results

### Wpl1p is necessary to mitigate Camptothecin and MMS induced cell cycle delay

Wpl1p has been implicated in regulating cohesin function in both cohesion and condensation. However, Wpl1p function in another cohesin-regulated process, DNA repair, has not been well characterized. In fact *WPL1* was originally characterized as a promoter of DNA repair, as *wpl1*Δ cells exhibited weak sensitivity to ionizing radiation (Game et al. 2003). Thus we were curious as to whether Wpl1p function in promotion of DNA repair was mediated through cohesin. We revisited the severe sensitivity of *eco1*Δ *wpl1*Δ cells to the topoisomerase I inhibitor, camptothecin (CPT) (Guacci and Koshland 2012). One likely cause of this CPT sensitivity was that *eco1*Δ caused a severe cohesion defect that impaired the use of the sister chromatid as an efficient template for DNA repair (Guacci and Koshland 2012). However, given Wpl1p’s role in promoting resistance to ionizing radiation, we wondered whether the loss of Wpl1p might contribute to the severe CPT sensitivity of *eco1*Δ *wpl1*Δ cells.

To assess the role of Wpl1p in CPT sensitivity, we compared the growth of wildtype, *wpl1*Δ and *eco1*Δ *wpl1*Δ cells on media containing 20 μg/mL CPT. As expected, *eco1*Δ *wpl1*Δ cells were unable to grow even after 5 days confirming that Eco1p plays a critical role in surviving DNA damage. Interestingly, *wpl1*Δ cells grew dramatically slower relative to wild-type cells, taking several additional days to form colonies, albeit smaller ones (Figure 1A). This growth delay suggests that Wpl1p may promote efficient DNA repair.

**Figure 1:**
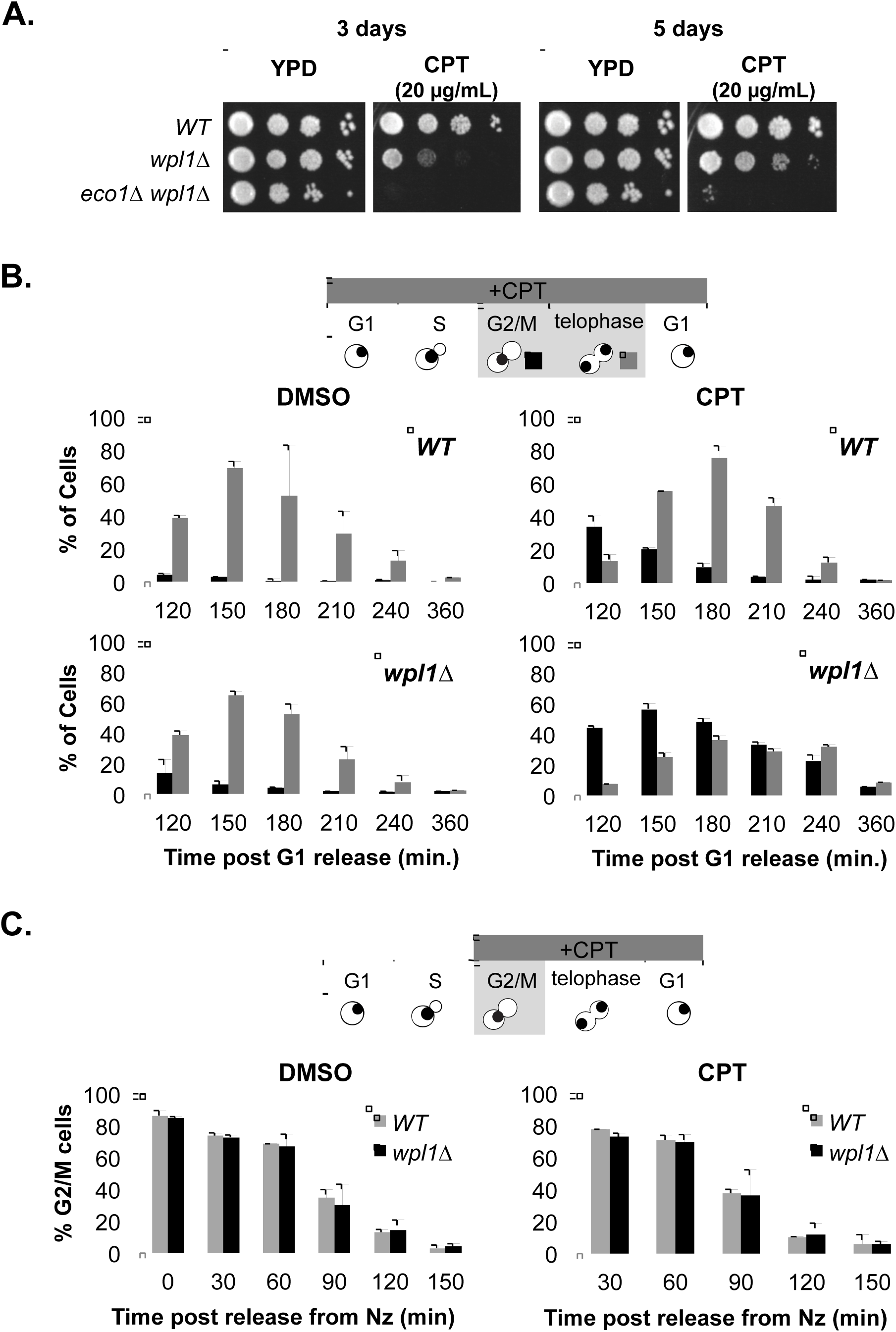
Wpl1p promotes efficient repair of camptothecin generated DNA damage. **(A)** *wpl1*Δ cells grow slowly on media containing camptothecin (CPT). *WT*(VG3349-1B), *wpl1*Δ (VG3360-3D), and *eco1*Δ *wpl1*Δ (VG3503 #4) cells were serially diluted (each spot represents 10x dilution), and plated on YPD media alone or containing 20 μg/mL CPT. Plates were incubated at 23°C and assessed at 3 and 5 days post plating. **(B)** *wpl1*Δ cells grown in the presence of camptothecin from G1 onward exhibit a prolonged mitotic delay. *WT*(VG3349-1B), and *wpl1*Δ (VG3360-3D) cells were grown tc mid-log phase in YPD at 23°C, arrested in G1 by addition of alpha-factor then released from G1 into YPD media buffered with 25 mM HEPES pH 7.4 as described in Materials and Methods. At the time of release cells were split into two aliquots and 20 μg/mL CPT was added to one and DMSO was added to the other. Once most cells had entered S-phase (90 minutes after release from G1), alpha-factor was added to ensure cells woulc progress through one cell-cycle and re-arrest in G1. Aliquots were taken every 30 minutes and fixed in 70% ethanol. Fixed cells were stained with DAPI to detect chromosomal DNA for scoring. Cells were scored for bud morphology (unbudded, small-medium bud, or large bud) and whether they contained a single DAPI chromosomal mass or two DAPI masses (top panel). Graphs show the percentage of large budded cells with a single DNA mass (G2/M; black) or two DNA masses (telophase; gray). **(C)** Camptothecin treatment of wild-type and *wpl1*Δ cells in mid-M phase has no effect on progression through mitosis. *WT* (VG3349-1B), and *wpl1*Δ (VG3360-3D) cells grown to mid-log phase in YPD at 23°C were arrested in G1 by addition of alpha-factor, then synchronously released from G1 into YPD containing nocodazole to re-arrest cells in mid-M phase. Cells were then released from mid-M arrest into YPD HEPES buffered media and cultures were split into two aliquots. 20 μg/mL CPT was added to one aliquot and DMSO to the other. Alpha factor was added to both aliquots to ensure cells exiting mitosis would arrest in G1. Aliquots taken every 30 minutes, fixed in 70% ethanol and then stained with DAPI and scored as described in B. Graphs show the percentage of large budded cells with a single DAPI mass (G2/M) at each time-point for WT cells (gray) or *wpl1*Δ cells (black)

To further characterize the kinetics of DNA damage repair, we compared how CPT treatment affected cell-cycle progression of wild-type and *wpl1*Δ cells. The extent and duration of a drug-induced cell-cycle delay serves as an indirect measure of DNA damage and repair. We synchronized wild-type and *wpl1*Δ cells in G1 with alpha-factor and then released the cells into media either containing CPT (20 μg/mL) or without CPT (DMSO) (Materials and Methods). Once cells had budded we reintroduced alpha-factor into the media to enable cells to progress through the cell cycle then re-arrest in the following G1. We collected aliquots of cells every 30 minutes after G1 release and analyzed them for bud morphology, DNA content, and chromosome segregation as measured by DNA morphology (Figure 1B).

Analysis of DNA content by flow-cytometry revealed that both wild-type and *wpl1*Δ cells treated with CPT progressed though S-phase (transitioning from 1C to 2C) with similar kinetics as did their DMSO-treated counterparts (Figure S1A). However, in CPT-treated *wpl1*Δ cells the 2C DNA peak persisted longer than in either CPT-treated wild-type cells or DMSO-treated cells of either genotype (Figure S1A; 180-240min). This result suggests that *wpl1*Δ cells delay in mitosis because of persisting CPT-generated damage that activated the G2/M DNA-damage checkpoint.

To provide more support for this putative G2/M delay, we examined the bud and DNA morphologies of these cells. During an unperturbed cell cycle in yeast, DNA replication is completed when the bud is small and a single nuclear DNA mass bearing all the chromosomes is present. As the cell cycle progresses, the bud grows to medium size and mitosis quickly ensues. As chromosomes segregate, two separated DNA masses of equal size can be distinguished, one in the mother cell and one in the bud (telophase cells). If cells stall prior to anaphase, the undivided nucleus remains at the bud neck while the bud continues to grow, giving rise large-budded cells with unsegregated chromosomes as seen by a single DNA mass (G2/M cells) (Figure 1B, top panel; (Hartwell 1974). As expected for control DMSO-treated wild-type and *wpl1*Δ cells, few large budded G2/M cells were observed, as most cells entered telophase when buds were mid-sized consistent with the absence of a cell-cycle delay (Figure 1B, left panels). By 150-minutes post release, most cells were in telophase (large-budded with divided nuclei). The number of telophase cells declined as cells underwent cytokinesis and by 240-minutes, most cells were arrested in G1 (Figure 1B left panels; Figure S1A).

When treated with CPT, both wild-type and *wpl1*Δ cultures exhibited a great increase in the amount of large-budded cells G2/M cells (~40% of cells) and few telophase cells were seen at 120-minutes post release compared to their DMSO-treated counterparts (Figure 1B, right panels). The similar G2/M cell cycle delays seen in both wild-type and *wpl1*Δ cells were consistent with both genotypes initially experiencing the same level of DNA damage when treated with CPT. Wild-type cells overcame this arrest quickly as seen by the high level of telophase cells at 150-minutes and 180-minutes, and that most cells exited mitosis by 240-minutes (Figure 1B, top-right panel). In contrast, the amount of *wpl1*Δ cells stalled in G2/M increased until 150-minutes with a significant amount remaining stalled through 240-minutes (Figure 1B, bottom-right panel). Eventually, all the stalled *wpl1*Δ cells entered telophase and exited mitosis, indicating that the CPT-induced damage was repaired. These results are consistent with a role for Wpl1p in the timely repair of CPT-induced damage.

CPT-mediated damage is thought to cause double-strand breaks (DSBs) when toposiomerase I-induced single-strand nicks encounter replication machinery during S-phase (Avemann et al. 1988; Strumberg et al. 2000; Saleh-Gohari et al. 2005). Thus, our results implicate Wpl1p as being important for the repair of S-phase induced damage. To test whether the delay observed in *wpl1*Δ cells was due to the DNA damage induced during S-phase, we allowed cultures of wild-type and *wpl1*Δ cells to progress synchronously through S-phase in the absence of CPT and arrest in G2/M by the addition of the microtubule poison, nocodazole. Cultures were released from nocodazole arrest and split in half and CPT was added to one aliquot while DMSO was added to the other. We then monitored progression every 30 minutes through mitosis and cytokinesis in either the presence or absence of CPT. Alpha-factor was added to the cultures to prevent progression of cells beyond the ensuing G1. Upon release from nocodazole, wild-type and *wpl1*Δ cells segregated their chromosomes with similar kinetics when treated with either DMSO or CPT (Figure 1C, Figure S1B). Thus, the CPT-induced delay in the initiation of chromosome segregation in *wpl1*Δ cells required the presence of CPT prior to M-phase. These results suggest that Wpl1p helps to repair CPT-induced damage during S-phase.

To address whether *WPL1* had a general role in mitigating other types of S-phase DNA damage, we analyzed the sensitivity of *wpl1*Δ cells to the alkylating agent, methyl-methane sulfonate (MMS). Consistent with our findings for CPT, sensitivity of *eco1*Δ *wpl1*Δ cells to MMS was previously reported, suggesting that the lack of cohesion in these cells is also detrimental for mitigation of MMS-induced damage (Sutani et al. 2009). We tested the sensitivity of *wpl1*Δ cells by monitoring their growth on media containing 0.01% MMS. The growth of *wpl1*Δ cells on MMS was significantly delayed compared to wild-type cells, taking several days to form colonies, similar to what was observed for CPT treatment (Figure 2A). The growth defect seen in these cells suggests that Wpl1p contributes positively to efficient repair of MMS-induced damage.

**Figure 2:**
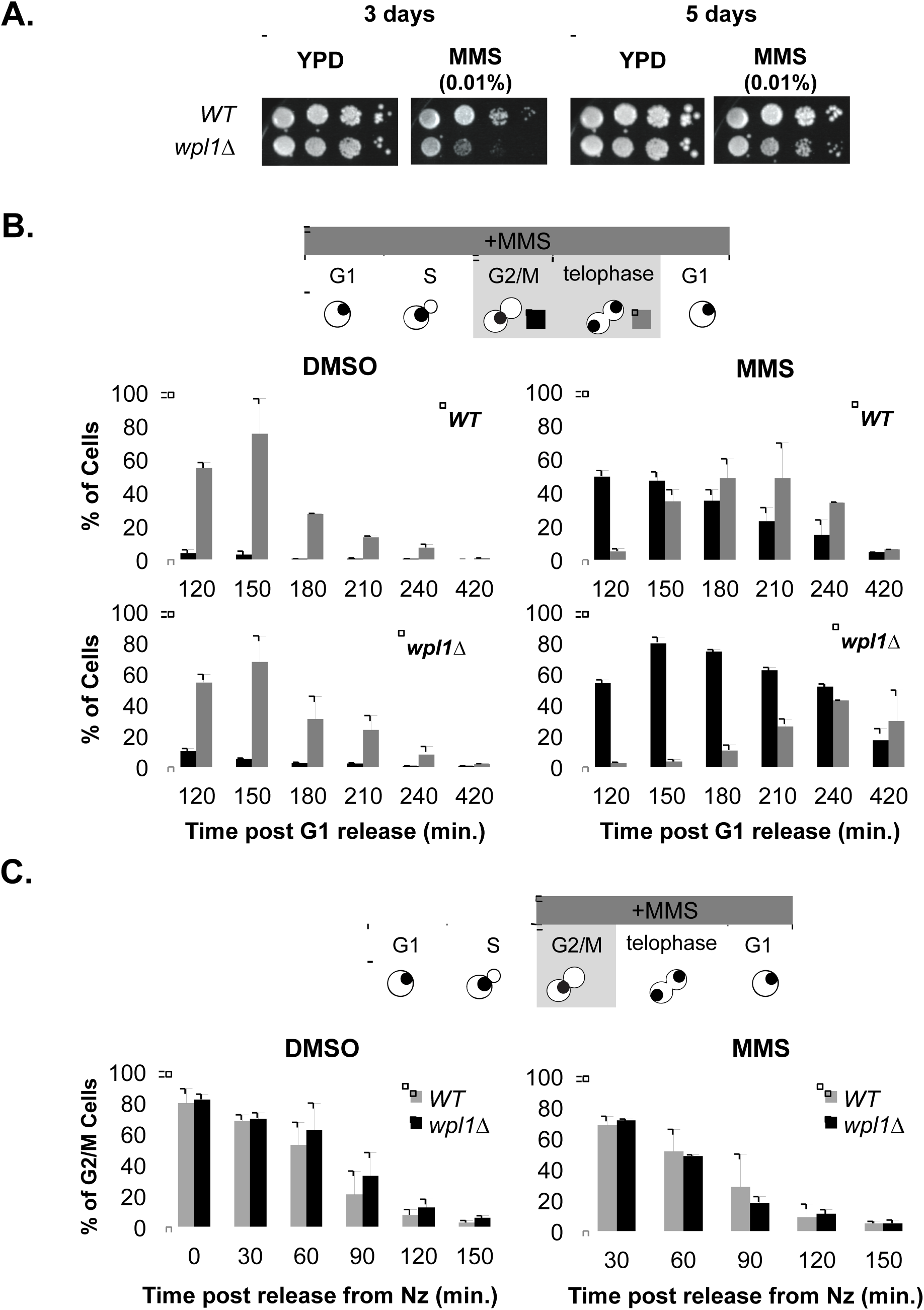
Wpl1p is necessary for efficient repair of MMS mediated DNA damage. **(A)** *wpl1*Δ cells are sensitive to MMS. Assessment of sensitivity of WT and *wpl1*Δ cells to MMS. *WT* (VG3349-1B) and *wpl1*Δ (VG3360-3D) cells were serially diluted 10x and plated onto YPD media either with or with out MMS to a final concentration of 0.01%. Plates were incubated at 23°C and assessed at 3 and 5 days post plating. **(B)** *wpl1*Δ cells grown in the presence of MMS from G1 onward exhibit a prolonged mitotic delay. *WT* (VG3349-1B) and *wpl1*Δ (VG3360-3D) cells synchronously released from G1 as described in Figure 1B, except MMS (0.01% final) was added instead of CPT and YPD media was not buffered. Cells were collected, processed, and scored as described in Figure 1B. Graphs show the percentage of large budded cells with a single DNA mass (G2/M; black) or two DNA masses (telophase; gray). **(C)** MMS treatment of wild-type and *wpl1*Δ cells in mid-M phase has no effect on progression through mitosis. *WT* (VG3349-1B) and *wpl1*Δ (VG3360-3D) cells were released from mid-M phase arrest as described in Figure 1C except cultures contained 0.01% MMS instead of CPT and YPD was not buffered. Graphs show the percentage of large budded cells with a single DAPI mass each time-point for *WT* cells (gray) or *wpl1*Δ cells (black).

We then tested the role of *WPL1* in mitigating MMS-damage in a single cell cycle by putting wild-type and *wpl1*Δ cells through the same regimen as described for CPT treatment. Treatment with 0.01% MMS did not slow the progression through S-phase for either wild-type or *wpl1*Δ cells but the 2C DNA peak persisted for both as compared to DMSO-treated cells (Figure S2A, 150-180min). This result indicated that MMS generated DNA damage that delayed cells in G2/M. However, the MMS-induced G2/M delay lasted much longer in *wpl1*Δ cells as they still showed a large 2C peak at 240-minutes whereas most wild-type cells had already entered G1 (1C peak) by 210 minutes (Figure S2A).

Analysis of bud and DNA morphologies in MMS-treated cells revealed that both wild-type and *wpl1*Δ cells experienced similar levels of damage-induced stalling in G2/M, as ~50% of cells of both genotypes were large-budded with undivided nuclei 120-minutes after release (Figure 2B right panels). However, by 180 minutes half of wild-type cells had already entered telophase whereas few *wpl1*Δ entered telophase so most remained stalled in G2/M. Moreover, ~20% of MMS-treated *wpl1*Δ cells remained in G2/M after 420-minutes suggesting that these cells were unable to recover at all. Finally, like CPT treatment, MMS addition to nocodazole-arrested cultures failed to induce a cell-cycle delay (Figure 2C, Figure S2B). Taken together, these results suggest that Wpl1p is important for the efficient repair of multiple types of DNA damage induced during S-phase.

### Destabilization of the Smc3p/Mcd1p interface promotes cohesion and DNA repair, and inhibits condensation

Several studies suggest that one molecular function of Wpl1p is to destabilize the interface between the N-terminus of Mcd1p and the base of the coiled-coil of Smc3p (Chan et al. 2012; Beckouët et al. 2016). We reasoned that Wpl1p’s destabilization activity might be important for promoting one or more of its regulatory functions of cohesin. If so, one or more of these regulatory functions would be compromised when Wpl1p’s destabilization activity was blocked by covalently fusing Smc3p to Mcd1p. Previously, a functional fusion of Smc3p fused to the N-terminus of Mcd1p was created (Gruber et al. 2006). We generated a strain in which the *SMC3*-*MCD1* fusion was the sole source of both *SMC3* and *MCD1* and assessed whether cells would phenotypically mimic a *wpl1*Δ. If so, the fusion strain should be defective for both efficient cohesion generation and DNA repair, as well as being unable to inhibit condensation.

We previously showed that restoration of viability to *eco1*Δ mutants requires the suppression of Wpl1p-dependent inhibition of condensation (Guacci and Koshland 2012). The Smc3-Mcd1p fusion was shown to restore viability to *eco1*Δ mutants (Chan et al. 2012), suggesting that the fusion may suppress Wpl1p’s ability to inhibit condensation. To assess this possibility directly, we examined chromosome condensation in an *SMC3*-*MCD1 eco1*Δ double mutant. We utilized a standard method for assessing yeast chromosome condensation by monitoring the repetitive *rDNA* locus (Guacci et al 1994; Guacci et al 1997). The *rDNA* locus is located in the nucleolus and protrudes from the bulk chromosomal mass making it easy to monitor its condensation state. A condensed *rDNA* locus forms a distinct loop structure, while decondensed *rDNA* locus form a “puff” morphology (Figure 3A; Materials and Methods) (Guacci et al. 1993; Guacci and Koshland 1994). We analyzed the morphology of the *rDNA* in cells that had been arrested in mid-M phase with nocodazole (Figure 3A). Most wild-type cells had condensed *rDNA* loops, with few displaying decondensed *rDNA* (Figure 3B). Using the auxin-inducible degradation system, we depleted Eco1p by the addition of auxin to cultures containing the *eco1*-*AID* allele and observed that over 80% of these cells had decondensed *rDNA.* Consistent with previous findings, only ~30% of *eco1*Δ *wpl1*Δ cells had decondensed *rDNAs*, indicating that the majority these cells were capable of mediating condensation. Similarly, only ~30% of *SMC3*-*MCD1 eco1*Δ cells had decondensed *rDNA* (Figure 3B). Thus, like *wpl1*Δ, the *SMC3*-*MCD1* fusion is able to restore condensation to cells lacking Eco1p.

**Figure 3:**
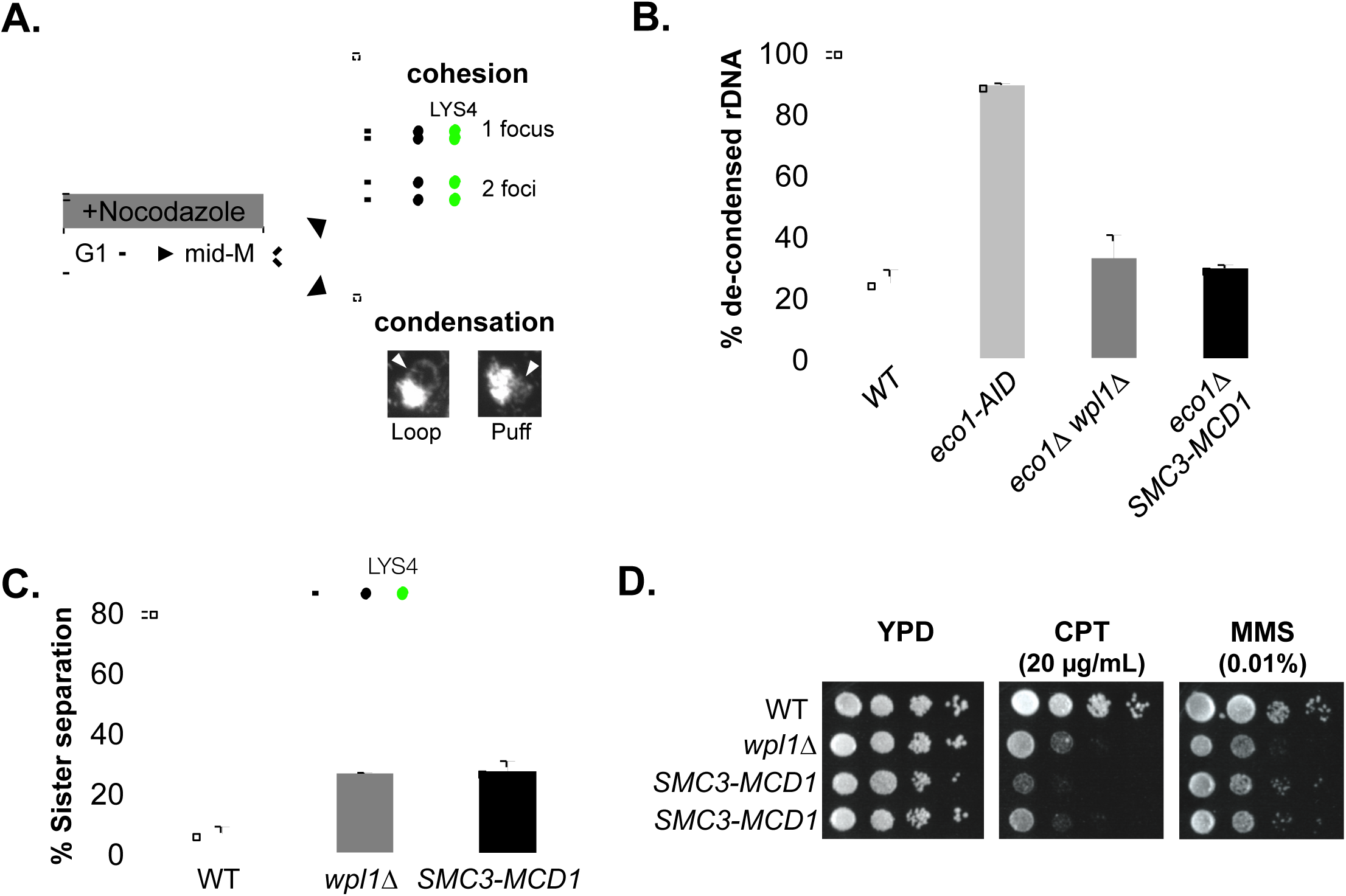
Smc3p/Mcd1p interface is important for cohesin function in cohesion, condensation and DNA damage. **(A)** Schematic of scoring cohesion and condensation. Cells were synchronously arrested in mid-M as described in Materials and Methods. Cells are processed for cohesion analysis of Lacl-GFP at *CEN*-distal *LYS4* locus and *CEN*-proximal *TRP1* and for condensation by FISH methodology as described in Materials and Methods. Chromosome condensation is assessed by morphology of the rDNA locus. Loop morphology indicates proper condensation while “puff” indicates a decondensed rDNA locus. **(B)** *SMC3*-*MCD1* fusion restores condensation in *eco1*Δ cells. *WT* (VG3349-1B), *eco1*-*AID (VG3633*-*2D)*, *eco1*Δ *wpl1*Δ (VG3502 #A) and *eco1*Δ *SMC3*-*MCD1* (MSB249-3A) were arrested in G1 using alpha factor then synchronously arrested in mid-M phase using nocodazole as described in material and methods. 500 μM auxin was present in the media of *eco1*-*AID* strain from G1 through mid-M phase. Cells were fixed and processed for FISH as described in Materials and Methods. rDNA condensation (loops) and defective condensation (puffs) were scored as described in Figure 3A. **(C)** *SMC3*-*MCD1* has similar modest cohesion defect to *wpl1*Δ. *PDS5* (VG3349-1B), *wpl1*Δ (VG3360-3D), and *SMC3*-*MCD1* (VG3940-2D) were arrested in mid-M phase using nocodazole as described in B. Cells were scored for cohesion (one GFP focus) and loss of cohesion (two GFP foci; sister separation) as described in A. The percentage of cells lacking cohesion (separation) is shown.

We next examined whether modulation of the Smc3p/Mcd1p interface was required for two other Wpl1p functions, promoting efficient cohesion and DNA damage repair. We tested whether the *SMC3*-*MCD1* fusion strain was as defective in efficiently promoting cohesion as a *wpl1*Δ strain. We monitored cohesion at a *CEN*-distal locus on chromosome IV by integration of LacO repeats at the *LYS4* locus in cells containing LacI-GFP (Materials and Methods). Cohesion was assessed in cells synchronously arrested in mid-M by treatment with nocodazole. Cells containing a single LacI-GFP focus indicated cohesion whereas cells containing two LacI-GFP foci indicated a loss of cohesion (Figure 3A). The *SMC3*-*MCD1* fusion strain exhibited a moderate cohesion defect of ~30%, similar to what is seen in a *wpl1*Δ strain (Figure 3C). Similar results for both the *SMC3*-*MCD1* fusion and the *wpl1*Δ strains were previously reported at the more CEN-proximal *URA3* locus (Gruber et al. 2006; Rowland et al. 2009).

To test whether the *SMC3*-*MCD1* fusion strain was also compromised for the ability to mitigate S-phase induced DNA damage, we compared wild-type, *wpl1*Δ and *SMC3*-*MCD1* fusion strains for sensitivity to 20 μg/mL CPT and to 0.01% MMS. The *SMC3*-*MCD1* fusion strain and the *wpl1*Δ strain showed similar growth inhibition to both drugs (Figure 3D). Thus, the Smc3-Mcd1p fusion protein mimics all the phenotypes of *wpl1*Δ for cohesion, condensation and sensitivity to DNA damage implying that blocking dissociation of Mcd1p from Smc3p is the common underlying molecular defect causing these defects. Taken together, these data suggest that Wpl1p destabilization of Smc3p/Mcd1p interface is necessary to promote cohesion and repair of S-phase induced DNA damage as well as to inhibit condensation.

### The Pds5p N-terminus regulates the promotion of cohesion and inhibition of condensation

Our findings along with previous studies suggest that Wpl1p regulation of diverse cohesin functions is complicated. To parse how Wpl1p distinguishes these functions, we sought to understand the relationship between Pds5p and Wpl1p. Given that Wpl1p and Pds5p form a complex, we wondered whether they cooperate to perform a common regulatory function. The potential for a functional cooperation is suggested by the fact that *wpl1*Δ and specific N-terminal *pds5* mutant alleles restore viability to *eco1*-*ts* cells that have reduced acetylase activity (Rowland et al. 2009; Chan et al. 2012). To further explore the functional relationship between Wpl1p and Pds5p, we asked whether *wpl1*Δ and three representatives of these N-terminal *pds5* alleles (*pds5*-*S81R*, *pds5*-*P89L*, and *pds5*-*E181K*) shared other signature phenotypes of *wpl1*Δ.

One signature phenotype of *wpl1*Δ is the restoration of viability to cells completely lacking Eco1p activity (*eco1*Δ) (Rowland et al. 2009; Sutani et al. 2009; Feytout et al. 2011). To test whether these *pds5* alleles could also suppress *eco1*Δ, *we* first constructed strains where *pds5*-*S81R*, *pds5*-*P89L*, or *pds5*-*E181K* was the sole *pds5* allele in cells. We then introduced a centromere plasmid containing *ECO1* and *URA3* (*ECO1 CEN URA3*) into these *pds5* mutant strains as well as into a wild-type *PDS5* strain. Finally, we deleted *ECO1* from its endogenous locus (*eco1*Δ) in all of these strains to generate *ECO1* shuffle strains. Counter-selection against cells containing the *ECO1 URA3 CEN* plasmid by plating on media containing 5-FOA will reveal whether any of the *pds5 eco1*Δ double mutants were viable.

As expected, cells containing wild-type *PDS5* (*WT*) and *eco1*Δ could not grow on 5-FOA as *ECO1* is an essential gene (Figure 4A). In contrast, both the *pds5*-*S81R* and *pds5*-*P89L* alleles enabled growth on 5-FOA indicating suppression of *eco1*Δ (Figure 4A). Consistent with our findings, previous results also showed that *pds5*-*P89L* restored viability to *eco1*Δ (Sutani et al. 2009). In contrast to *pds5*-*S81R* and *pds5*-*P89L*, *pds5*-*E181K* was unable to support viability to *eco1*Δ (Figure 4A). To determine whether the inability of *pds5*-*E181K* to restore viability to *eco1*Δ was due to weak suppressor activity, we rebuilt *pds5*-*E181K* into a strain containing the *eco1*-*203* temperature-sensitive allele. At the restrictive temperature, 34°C, *pds5*-*E181K* was able to restore viability to *eco1*-*203* cells (Figure S3A). This result is consistent with a previous report in which *pds5*-*E181K* suppressed the inviability of the *eco1*-*1* temperature sensitive allele (Rowland et al. 2009). Thus, *pds5*-*S81R* and *pds5*-*P89L* are akin to a *wpl1*Δ as they suppress an *eco1*Δ whereas *pds5*-*E181K* can only suppress inviability when Eco1p function is reduced but not abolished.

**Figure 4:**
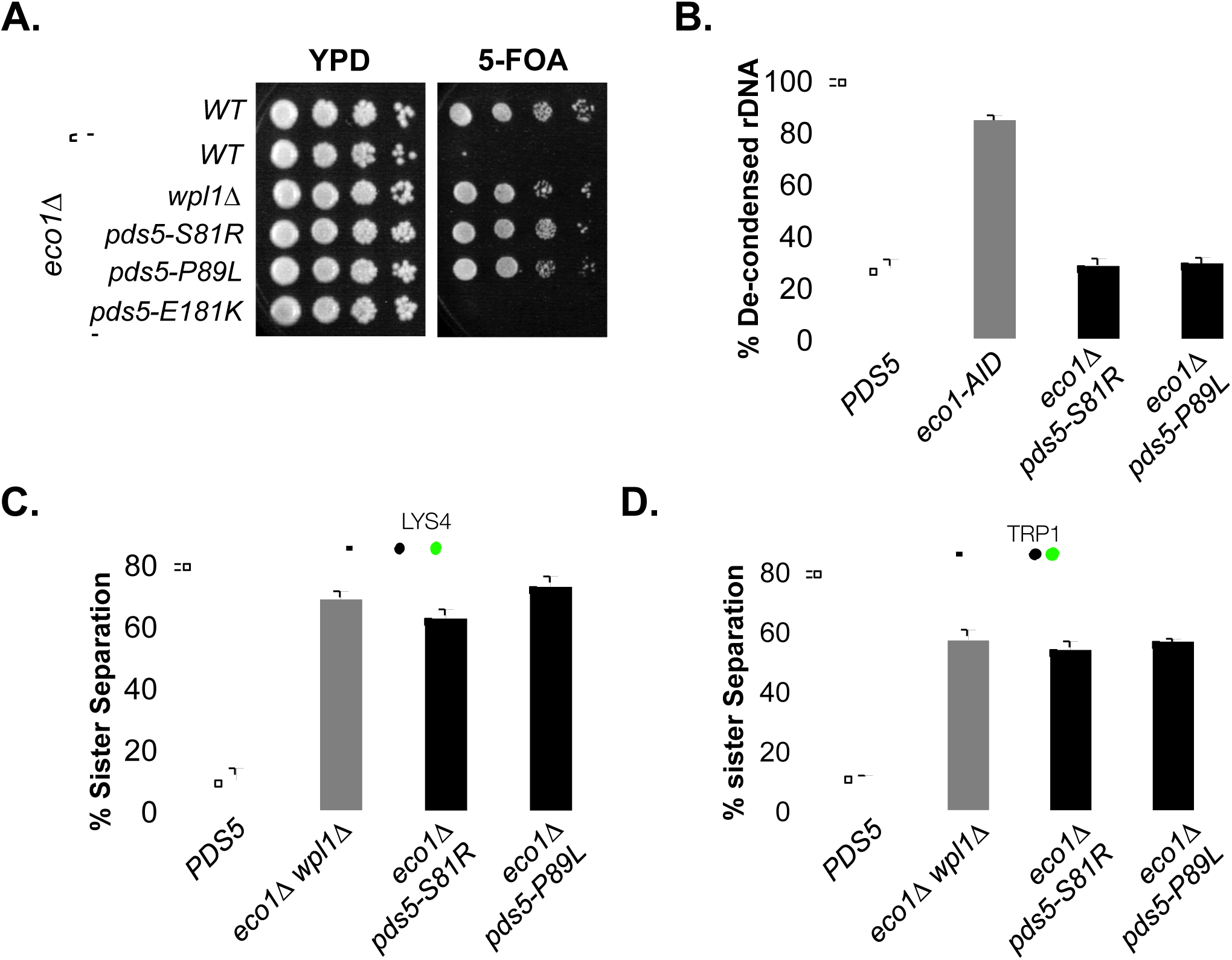
*pds5* N-terminal mutants suppress inviability of *eco1*Δ through restoration of condensation. **(A)** Plasmid shuffle assay to assess viability of *pds5* N-terminal mutants in *eco*Δ background. Plasmid pBS1030 *(ECO1 CEN URA3)* is present in haploid wild-type (VG3349-1B), *eco1*Δ (VG3499-1B), *eco1*Δ wpl1Δ (VG3503 #4), *eco1*Δ *pds5*-*S81R* (MSB138-1K), *eco1*Δ *pds5*-*P89L* (MSB139-2J), *eco1*Δ *pds5*-*E181K* (MSB147-1A) strains. Cells were grown in YPD media then plated at 10x dilution on YPD or 5-FOA media at 23°C for 3 days. 5-FOA selects for loss of pBS1030 (*ECO1 CEN URA3*). **(B)** *pds5*-*S81R* and *pds5*-*P89L* restore condensation in *eco1*Δ cells. *PDS5* (VG3349-1B), *eco1*-*AID (VG3633*-*2D)*, *eco1*Δ *pds5*-*S81R* (MSB138-1K), *and eco1*Δ *pds5*-*P89L* (MSB139-2J) were synchronously arrested in mid-M phase as described in B except that 500 μM auxin was present in the media of *eco1*-*AID* strain from G1 through mid-M phase. Cells were fixed and processed for FISH as described in Materials and Methods. *rDNA* condensation (loops) and defective condensation (puffs) were scored as described in Figure 3A. **(C&D)** *pds5*-*S81R eco1*Δ and *pds5*-*P89L eco1*Δ double mutants have a dramatic defect in cohesion. *PDS5* (VG3349-1B & MSB185-1A), *eco1*Δ *wpl1*Δ (VG3503 #4 & VG3502 #A), *eco1*Δ *pds5*-*S81R* (MSB138-1K & MSB210-2A), *eco1*Δ *pds5*-*P89L* (MSB139-2J & MSB211-2J) were arrested in G1 using alpha factor then synchronously arrested in mid-M phase using nocodazole as described in material and methods. Cells were scored for cohesion (one GFP focus) and loss of cohesion (two GFP foci; sister separation) as described in Figure 3C. The percentage of cells lacking cohesion (separation) is shown.

A second signature phenotype of *wpl1*Δ js the restoration of condensation but not cohesion to *eco1*Δ cells (Guacci and Koshland 2012). This pattern of suppression distinguishes inactivation of Wpl1p function from other suppressors of *eco1*Δ in the cohesin ATPase that restore both cohesion and condensation (Guacci and Koshland 2012; Çamdere et al. 2015). To test whether these *pds5* N-terminal alleles phenocopied this *wpl1*Δ phenotype, we assessed the chromosome condensation in *pds5*-*S81R eco1*Δ and *pds5*-*P89L eco1*Δ cells arrested in mid-M phase (Figure 3A). Most wild-type (*PDS5*) cells arrested in mid-M had condensed *rDNA* loops so few had decondensed *rDNA* (Figure 4B). We depleted Eco1p by the addition of auxin to cultures containing the *eco1*-*AID* allele and observed that over 80% of these cells had decondensed *rDNA.* Similar to what was previously reported for *eco1*Δ *wpl1*Δ cells, most *pds5*-*S81R eco1*Δ and *pds5*-*P89L eco1*Δ cells had condensed *rDNA* loops so only ~20-30% of cells exhibited decondensed rDNA loci (Figure 4B) (Guacci and Koshland 2012).

Through a similar regimen, we assessed cohesion in *pds5*-*S81R eco1*Δ and *pds5*-*P89L eco1*Δ double mutant cells by monitoring cohesion of mid-M phase arrested cells at *CEN*-proximal (*TRP1*) and *CEN*-distal (*LYS4*) loci on chromosome IV (Figure 3A). As expected from our previous studies (Guacci and Koshland 2012), most sister chromatids remained tethered in both regions in wild-type cells whereas ~70% of *eco1*Δ *wpl1*Δ cells exhibited separated sister chromatids (Figure 4C & D). Similar to *eco1*Δ *wpl1*Δ cells, both *pds5*-*S81R eco1*Δ and *pds5*-*P89L eco1*Δ cells also exhibited high levels of separated sisters at both *CEN*-proximal and distal loci (Figure 4C & D). Additionally, the *pds5*-*E181K eco1*-*203* double mutant and *eco1*-*203* single mutant strains also had high levels of sister separation similar to *eco1*Δ *wpl1*Δ cells (Figure S3B). Thus, the *pds5* N-terminal suppressor mutants behave like *wpl1*Δ as they restore viability and condensation but not cohesion to *eco1*Δ or *eco1*-*ts* cells.

The third signature phenotype of *wpl1*Δ cells is its partial cohesion defect, ~30% (Figure 3C) (Guacci and Koshland 2012). Pds5p defective cells were already known have a severe cohesion defect (~80%) (Hartman et al. 2000; Stead et al. 2003; Noble et al. 2006; Guacci and Koshland 2012; Tong and Skibbens 2014). This quantitative difference suggested that Wpl1p and Pds5p might promote cohesion by different mechanisms. Alternatively, Pds5p might promote cohesion by two mechanisms, one dependent on Wpl1p and the other independent of Wpl1p. Given the phenotypic similarity between *wpl1*Δ and the N-terminal alleles of *pds5* described above, we wondered whether the N-terminus of Pds5p might be involved in a Wpl1p-dependent pathway for cohesion. To test this, we monitored the ability of the *pds5*-*S81R*, *pds5*-*P89L*, and *pds5*-*E181K* to mediate cohesion either in the presence or in the absence of *WPL1.*

We first assessed cohesion in the *pds5* N-terminal mutants in an otherwise wild-type background (*WPL1*) in cells arrested at mid-M phase using nocodazole. When cohesion was monitored at both *CEN*-proximal and *CEN*-distal loci, all three *pds5* N-terminal mutants exhibited cohesion defects of ~20% and ~30%, respectively, similar to that of *wpl1*Δ (Figure 5A & B). Additionally, kinetic analysis of the *pds5* N-terminal mutants showed that they lost cohesion similarly to *wpl1*Δ as cells progressed from S-phase to M-phase (Figure S4). These quantitative similarities were consistent with the model that Wpl1p and the Pds5p N-terminal domain acted in a common pathway to promote efficient cohesion establishment.

**Figure 5:**
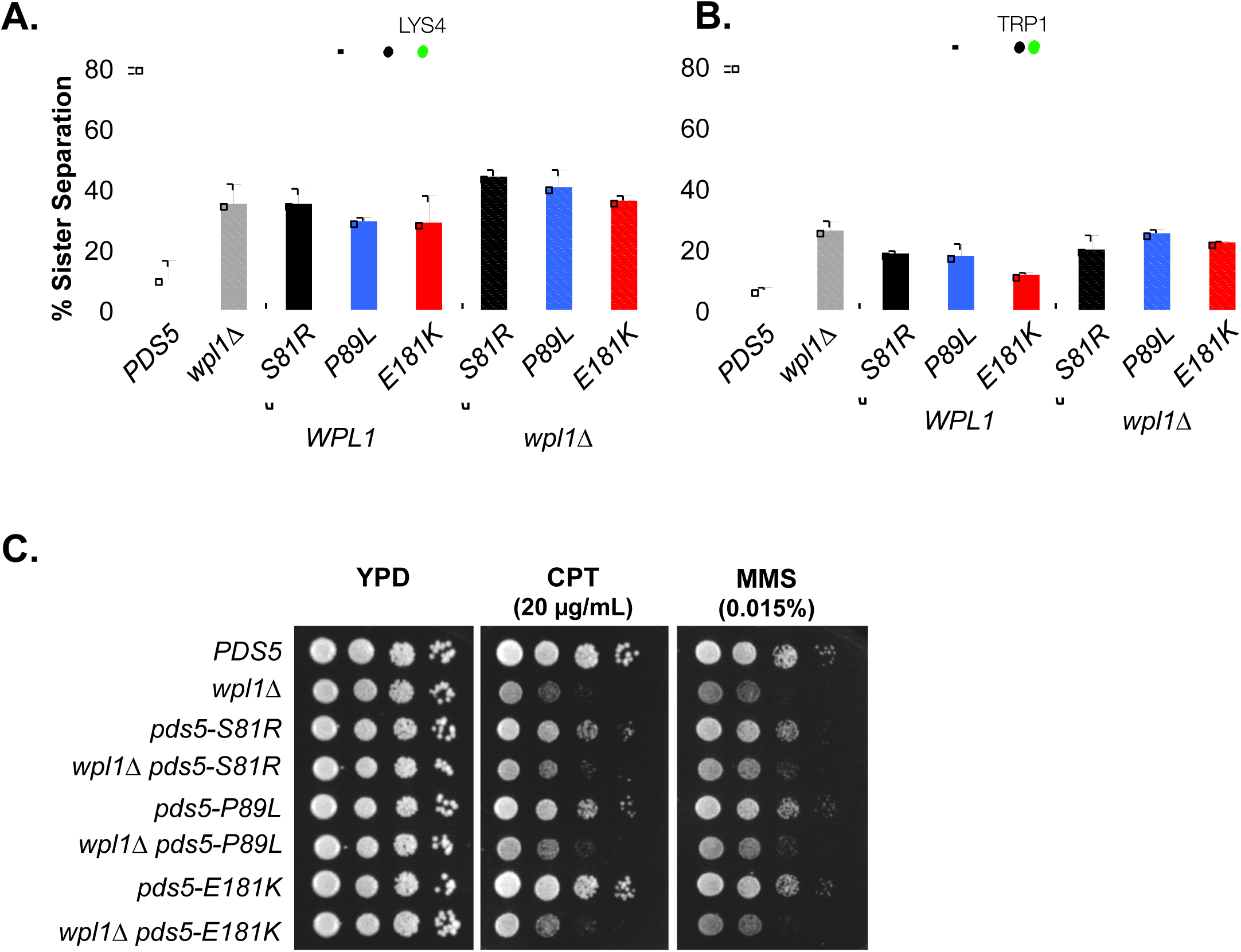
*pds5* N-terminal mutants are defective for Wpl1p-mediated cohesion, but not DNA repair. **(A&B)** *pds5* N-terminal mutants have similar modest cohesion defect as *wpl1*Δ alone or as *pds5 wpl1*Δ double mutants. Cells were synchronously arrested in mid-M as described in Materials and Methods and scored for cohesion at the *CEN*-distal *LYS4* and *CEN*-proximal *TRP1* locus as described in Figure 4A. Strains *PDS5* (VG3349-1B & MSB185-1A), *wpl1*Δ (VG3360-3D & VG3513-1B), *pds5*-*S81R* (MSB183-1A & MSB190-3E), *pds5*-*P89L* (MSB184-3A & MSB191-3A), *pds5*-*E181K* (MSB101-3C & MSB186-2E), *pds5*-*S81R wpl1A* (MSB133-3C & MSB204-1B), *pds5*-*P89L wpl1A* (MSB134-1L & MSB205-4C), *pds5*-*E181K wpl1A* (MSB223-1A & MSB206-6A) were assessed for cohesion loss (sister separation) and plotted. **(C)** Assessment of sensitivity of *pds5* N-terminal mutants to CPT and MMS. Cultures of cells in *WPL1* background: *PDS5* (VG3349-1B) *pds5*-*S81R* (MSB183-1A), *pds5*-*P89L* (MSB184-3A), and *pds5*-*E181K* (MSB101-3C), and *wpl1*Δ background: *wpl1*Δ (VG3360-3B), *pds5*-*S81R wpl1*Δ (MSB204-1B), *pds5*-Δ *P89L wpl1*Δ (MSB205-4C), *pds5*-*E181K wpl1*Δ (MSB223-1A), were serially diluted and plated on YPD media either containing no drug, 20 μg/mL CPT or 0.015% MMS, and incubated at 23°C and assessed at 3 days post plating.

To determine whether Wpl1p and the Pds5p N-terminus function in a common pathway in promoting cohesion, we assessed whether *wpl1*Δ enhanced the cohesion defect of the *pds5*-*S81R*, *pds5*-*P89L and pds5*-*E181K.* The cohesion defects of all three *pds5 wpl1*Δ double mutants in mid-M arrested cells were the same or only slightly higher than *pds5* single mutants alone or *wpl1*Δ alone (Figure 5A & B). If the *pds5* mutants and *wpl1*Δ promoted cohesion by distinct mechanisms, we would expect an additive effect in the double mutants, and would expect to see cohesion loss approaching 70% in these cells. The fact that *pds5* single mutants and *pds5 wpl1*Δ double mutants have similar partial defects in cohesion suggests that the Pds5p N-terminus and Wpl1p promote cohesion through a common pathway. These results suggest that Wpl1p interacts functionally with Pds5p both to inhibit condensation and to efficiently promote cohesion.

The final signature *wpl1*Δ phenotype is the sensitivity to the DNA damaging agents, CPT and MMS. As our results suggest that the N-terminus of Pds5p and Wpl1p function together in cohesion and condensation, we wondered whether they also function together to promote DNA damage repair. To test this possibility, we examined effects on the growth of the *pds5* N-terminal mutants alone or in the *wpl1*Δ background by plating the single and double mutants on media containing either CPT or MMS. *pds5*-*S81R*, *pds5*-*P89L* and *pds5*-*E181K* alone grew similarly to wild-type on 20 μg/ml CPT and 0.015% MMS and significantly better than *wpl1*Δ cells. Furthermore, *wpl1*Δ *pds5* double mutants and *wpl1*Δ cells were equally sensitive to both CPT and MMS (Figure 5C). These results suggest that Wpl1p’s role in DNA damage repair is independent of its functional interaction with the Pds5p N-terminus.

### Wpl1p interaction with Pds5p N-terminus is not sufficient for regulation of cohesin function

A simple model for a common biological function of Wpl1p and the N-terminus of Pds5p in cohesion and condensation is that these functions derive from the Wpl1p-Pds5p complex. If so, *pds5* N-terminal mutants might inhibit the formation of the Wpl1p-Pds5p complex. Support for this idea came from a recent crystal structure of human Pds5B (Ouyang et al. 2016). This crystal structure also contained a short peptide of Wapl, the human ortholog of Wpl1p, which bound to the N-terminus of Pds5B. As this region of Pds5B was highly conserved with yeast Pds5p, we were able to map the analogous residues of the Pds5p N-terminal mutations on the crystal structure of Pds5B (Figure 6A, Figure S5). These residues were located either within or in very close proximity to the Wapl binding site (Figure 6A). Additionally, yeast Wpl1p contains a partial consensus sequence to the conserved [K/R] [S/T] YSR motif important for Wapl interaction with Pds5B in vertebrates, suggesting that Wpl1p and Pds5p may bind in a similar manner in yeast (Ouyang et al. 2016).

**Figure 6:**
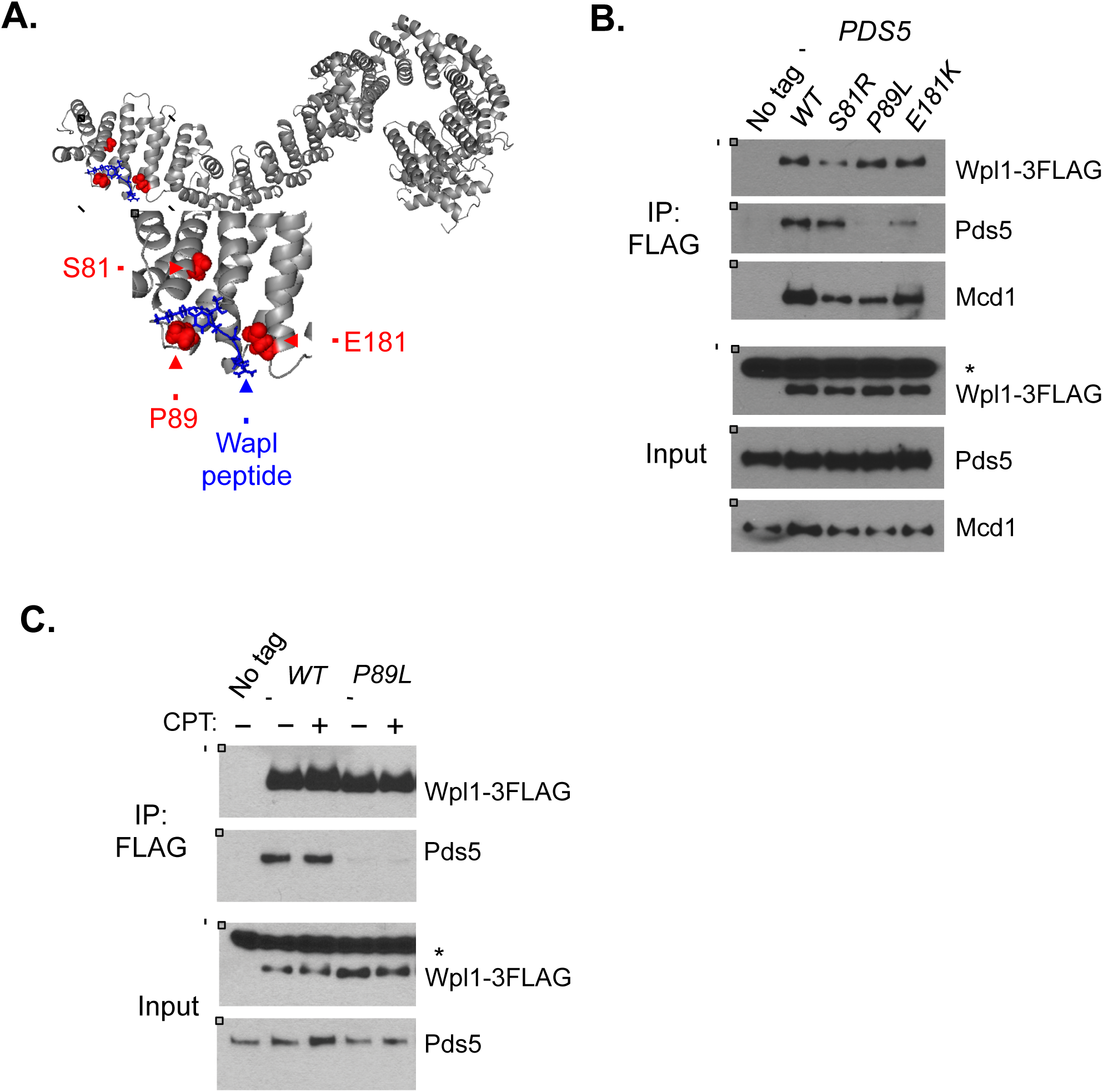
Pds5p N-terminus promotes Wpl1p binding to cohesin complex. **(A)** Crystal structure of Pds5B bound to YSR motif of Wapl from (Ouyang et al. 2016). Gray: Pds5B Blue: Wapl peptide Red: *eco1*-*ts* suppressors from (Rowland et al. 2009) and (Sutani et al. 2009). Yeast residues were mapped to analogous residues on Pds5B through alignment. **(B)** Pds5p N-terminal mutants impair Wpl1p binding to cohesin and can impair Wpl1p interaction with Pds5p. Wpl1-3FLAG was immunoprecipitated from protein extracts in asynchronous cultures containing Wpl1p-3FLAG and either *PDS5* (MSB192-2A), *pds5*-*S81R* (MSB193-1B), *pds5*-*P89L* (MSB194-1C), or *pds5*-*E181K* (MSB195-2D) as described in materials and methods. “No tag” control contains wildtype untagged *WPL1* and *PDS5* alleles (VG3349-1B). For western blot analysis, Wpl1p was detected using mouse anti-FLAG, Pds5p was detected using rabbit anti-Pds5, and Mcd1p was detected using rabbit anti-Mcd1 antibodies (Materials and Methods). For anti-FLAG, a non-specific species present in all cells is denoted by asterisk. **(C)** Assessment of interaction between Wpl1p-3FLAG and Pds5p (MSB192-2A) or Pds5p-P89L (MSB194-1C) when treated with CPT. Asynchronous cultures were treated either with DMSO or 20 μg/mL CPT for 3 hours before being harvested. No tag control is *WPL1 PDS5* (VG3349-1B) strain that was treated with DMSO. Immunoprecipitation and western blot were performed as described in part B and Materials and Methods.

Given this structural information, we asked whether the Pds5p N-terminal mutations disrupted the physical interaction between of Pds5p and Wpl1p. We C-terminally tagged Wpl1p with the Flag epitope (Wpl1p-3FLAG) and then performed anti-FLAG immunoprecipitation from extracts of asynchronously growing cells. We compared Wpl1p co-immunoprecipitation with wild-type Pds5p and each of the N-terminal Pds5p mutants. Anti-FLAG immunoprecipitation robustly co-immunoprecipitated wild-type Pds5p when Wpl1p-3FLAG was present but not when Wpl1p was untagged, confirming that Pds5p co-immunoprecipitation is due to a specific interaction with Wpl1p (Figure 6B). The Wpl1p-3FLAG immunoprecipitates contained very little Pds5p-P89L and significantly reduced Pds5p-E181K (Figure 6B). Thus, both mutations disrupt Pds5p binding to Wpl1p. In contrast, Pds5p-S81R retained binding to Wpl1p-3FLAG at a level similar to wild-type Pds5p (Figure 6B). These differences in Wpl1p binding between the three N-terminal Pds5p mutants are surprising as all three mutants similarly disrupt promotion of cohesion, and can restore viability to *eco1* mutants through restoration of condensation. These results suggest that binding to the Pds5p N-terminus is required for Wpl1p’s function as both an inhibitor of condensation and efficient promoter of cohesion. However, this interaction is not sufficient for these functions, as Pds5p-S81R bind Wpl1p-3FLAG at close to wild-type levels.

Despite the differences in binding between Wpl1p and Pds5p among the three *pds5* mutants, they have similar functional defects *in vivo.* These results suggest that the molecular function of Wpl1p must be attenuated through a mechanism other than Pds5p binding. Thus, we wondered whether the interaction between Wpl1p and cohesin might be compromised in these mutants. We re-probed the Wpl1p-3FLAG immunoprecipitates for the cohesin subunit, Mcd1p. In the wild-type Pds5p strain, Mcd1p exhibited a robust co-immunoprecipitation with Wpl1p (Figure 6B). In contrast, in all three Pds5p N-terminal mutant strains, there was reduced Mcd1p co-immunoprecipitation with Wpl1p (Figure 6B). Thus, we conclude that formation of a functional Wpl1p-Pds5p complex is important for efficient recruitment of Wpl1p to cohesin. Additionally, Wpl1p was still able to interact with Mcd1p in *pds5*-*P89L* mutant cells despite the Wpl1p interaction with Pds5p being abolished. This result indicates that Wpl1p can bind cohesin independently of Pds5p, which corroborates previous studies in yeast and other organisms that show that Wpl1p can interact directly with the cohesin subunit Scc3p/SA/STAG (Rowland et al. 2009; Shintomi and Hirano 2009).

In contrast to our conclusion that the Pds5p N-terminus functions with Wpl1p to inhibit condensation and promote cohesion, our studies suggest that Wpl1p can function independently of the Pds5p N-terminus in promotion of DNA repair. Consistent with this idea, our results show that *pds5*-*P89L* abrogates Wpl1p interaction with Pds5p but Wpl1p retains binding to Mcd1p. However, it is possible that DNA damage causes a modification to Pds5p or Wpl1p that promotes formation of the Wpl1p-Pds5p complex. If so, the interaction between Wpl1p and Pds5p-P89L might be restored upon induction of DNA damage. To assess this possibility, we treated asynchronously growing *PDS5 WPL1*-*3FLAG* and *pds5*-*P89L WPL1*-*3FLAG* cells with either DMSO or 20 μg/mL CPT for 3 hours. We immunoprecipitated Wpl1p from extracts from these cells and assayed for Pds5p binding. Wild-type Pds5p and Wpl1p co-immunoprecipitated at similar levels with or without CPT whereas Pds5p-P89L remained unable to co-immunoprecipitate with Wpl1p under either condition (Figure 6C). These findings further corroborate our conclusion that Wpl1p promotes DNA damage repair independently of its interaction with Pds5p.

## Discussion

Previous studies in budding yeast had demonstrated roles for Wpl1p in promoting efficient sister chromatid cohesion and in inhibiting condensation (Guacci and Koshland 2012; Lopez-Serra et al 2013). Here we provide evidence for a biological function of Wpl1p in the timely repair of DNA damage in S-phase, beyond its roles in cohesion and condensation. We report that cells blocked for Wpl1p function grow slowly when they experience DNA damage induced during S-phase by CPT and MMS. This slow growth results from a delay in the onset of chromosome segregation, likely reflecting activation of the DNA damage checkpoint because of slow repair of the damage.

The defect in DNA repair in cells blocked for Wpl1p function cannot be explained by their partial cohesion defect. We showed that cells containing the *pds5* mutants have the same partial cohesion defect as *wpl1*Δ and *SMC3*-*MCD1* cells but are phenotypically identical to wild-type cells when treated with CPT or MMS. These results suggest that Wpl1p modulates a cohesin function in the repair of S-phase-induced DNA damage beyond simply its role in promoting sister chromatid cohesion. When DNA damage is induced in G2/M, cohesin loading around the break-site is stimulated (Ström et al. 2004; Unal et al. 2004). Thus, it is possible that this positive role of Wpl1p is to promote cohesin binding at either sites of DNA damage or at replication forks to reinforce them upon damage.

The results from our study suggest that Wpl1p regulates cohesin function in DNA repair, cohesion and condensation through a common mechanism. We show that cells expressing the Smc3p-Mcd1p fusion protein, like *wpl1*Δ, show partial cohesion defects and growth sensitivities to S-phase DNA damaging agents, and restore viability and condensation to cells lacking Eco1p. Wpl1p is known to destabilize the interface between Smc3p and Mcd1p. The fusion of Smc3p and Mcd1p makes this interface refractory to Wpl1p function and like *wpl1*Δ, hyper-stabilizes the interaction between Mcd1p and Smc3p (Beckouët et al. 2016). Thus the destabilization of the Smc3p/Mcd1p interface, presumably through Wpl1p function, is required for efficient cohesion, timely repair of DNA damage and inhibition of condensation.

Disruption of the Smc3p/Mcd1p interface is thought to be one mechanism to unload cohesin from DNA (Chan et al. 2012). The results of our study suggest that Wpl1p-mediated unloading of cohesin has both positive (efficient promotion of cohesion and DNA repair) and negative (inhibition of condensation) consequences. Previous work suggested that cohesin and Pds5p regulate condensation by first binding DNA at sites along a chromatid then interactions between cohesins bound along the same chromatid (in *cis*) are generated to loop out the intervening DNA to generate axial shortening (Guacci et al 1997; Hartman et al 2000). The negative impact of Wpl1p on condensation is straightforward. Wpl1p either destabilizes the interaction between non-acetylated cohesin and/or destabilizes DNA binding of non-acetylated cohesins. This Wpl1p activity thereby prevents cohesin from tethering DNA in *cis* along a chromatid to inhibit condensation (Figure 7A). It is curious, though, that destabilization of cohesin’s interaction with DNA, would promote DNA repair and cohesion. These positive aspects may reflect cohesin’s burden to carry out diverse biological functions. In addition to loading cohesin prior to S-phase to establish cohesion, cohesin is known to be loaded *de*-*novo* at sites of damage and at stalled replication forks (Ström et al. 2004; Unal et al. 2004; Tittel-Elmer et al. 2012). These spatial and temporal constraints may require a cohesin pool that can be mobilized either from the nucleoplasm or from DNA binding at non-productive sites in the genome (Figure 7B top). In the absence of Wpl1p function, cohesin is stabilized on DNA so is less efficiently mobilized, and perhaps trapped in such non-productive sites. Additionally, in *wpl1*Δ cells cohesin levels on DNA and in cells are decreased (Sutani et al. 2009; Guacci et al. 2015) These effects would thereby limit the pool of dynamic cohesin. Thus cohesion promotion is less efficient both during a normal cell cycle and in response to DNA damage (Figure 7 bottom).

**Figure 7:**
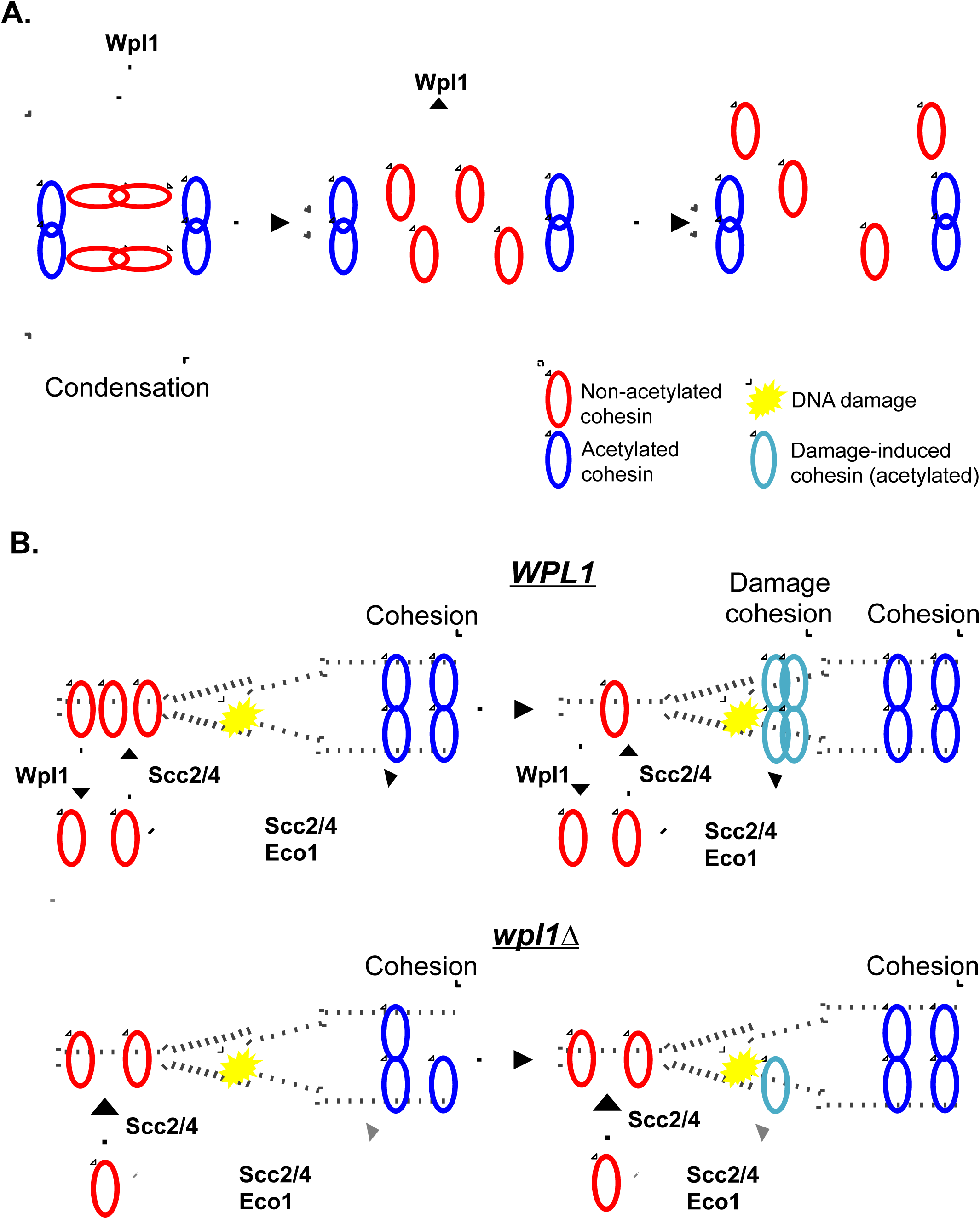
Model for Wpl1p promotion and inhibition of cohesin function through recycling. **(A)** Wpl1p inhibits condensation. Cohesin mediates condensation through chromosome looping. Non-acetylated cohesin (red) promotes condensation, while acetylated cohesin (blue) promotes cohesion. Wpl1p antagonizes condensation by removing non-acetylated cohesin from DNA. **(B)** Wpl1p promotes cohesion and DNA damage repair **Top:** non-acetylated cohesin (red) is loaded onto DNA by Scc2/4p, and is removed from DNA by Wpl1p, maintaining a soluble pool of cohesin. Cohesin loading, followed by Eco1p acetylation promotes cohesion, which is refractory from Wpl1p. Upon DNA damage, non-acetylated cohesin is removed from other sites in the genome and loaded around damage-site. **Bottom:** In the absence of Wpl1p, cohesin is loaded onto DNA by Scc2/4p but cannot be removed, causing the soluble pool of cohesin to be smaller. Thus, cohesion establishment is less efficient. Upon DNA damage, cohesin cannot be removed from other sites in the genome. Thus, cohesin loading around damage-site is less efficient.

The necessity of maintaining a dynamic pool of cohesin is supported by a number of observations. First, only ~20-30% of cohesin is acetylated during S-phase to establish sister chromatid cohesion (Zhang et al. 2008). As acetylated cohesin is thought to be refractory to Wpl1p activity (Rolef Ben-Shahar et al. 2008; Unal et al. 2008), this low level of acetylation may ensure that a substantial portion of cohesin remains responsive to Wpl1p. Second, we previously showed that a reduction in the total cellular pool of cohesin leads to more severe defects in condensation and DNA repair than cohesion (Heidinger-Pauli et al. 2010). These phenotypes may arise from a larger proportion of the remaining cohesin being locked onto the DNA in the cohesive (acetylated) state that is refractory to Wpl1p and thus not available for recycling in order to promote condensation and DNA repair. In this light, the primary biological function of Wpl1p is not to inhibit cohesin function by removing it from DNA. Rather, it would be to generate a dynamic cohesin pool for re-distribution to different chromosomal sites to perform cohesin’s distinct biological functions.

The idea that pools of cohesin need to be mobilized by Wpl1p to enable cohesin to perform different biological functions can explain two seeming paradoxes from our studies. The first paradox is the finding that the three *pds5* N-terminal mutants dramatically differ in their ability to bind Wpl1p, yet they phenocopy a *wpl1*Δ in their failure to efficiently promote cohesion and inhibit condensation. The second paradox is that the three *pds5* mutants differ from a *wpl1*Δ in that they are not sensitive to DNA damaging agents. All three *pds5* N-terminal mutants do share a common molecular defect: a reduction in the amount of Wpl1p bound to cohesin. This finding can explain the twin paradoxes described above. The regulation of both cohesion and condensation entail modulation of cohesin at many sites genome-wide. *pds5* N-terminal mutants may reduce the amount of Wpl1p bound to cohesin below a threshold required to mobilize the global pool of cohesin and thus impair proper regulation of cohesion and condensation. In contrast, DNA damage repair should only involve cohesin at a limited number of sites within the genome. This small requirement to mobilize enough cohesin to promote DNA repair may be met by the reduced level of Wpl1p in the *pds5* N-terminal mutants. As there is no Wpl1p present in *wpl1*Δ cells, all three cohesin biological functions would be defective.

In addition to the insights into the relationship between Wpl1p and cohesin, our work also furthers our understanding of the relationship between Wpl1p and Pds5p. Two *pds5* N-terminal alleles either entirely (*pds5*-*P89L*) or partially (*pds5-E181K*) disrupt the interaction with Wpl1p. However, in these cells, Wpl1p still binds cohesin, albeit at reduced levels. These data suggest that one function of the Wpl1p-Pds5p complex is to help recruit Wpl1p to cohesin. However, Wpl1p can also bind cohesin independent of its ability to complex with Pds5p. This independence has previously been demonstrated *in vitro* (Rowland et al. 2009; Shintomi and Hirano 2009). The *pds5*-*S81R* mutation preserves the interaction between Pds5p and Wpl1p, yet its effects on cohesin function are the same as *pds5*-*P89L*, which abolishes this interaction. This result indicates that formation of a Wpl1p-Pds5p complex is not sufficient for Wpl1p function. It may be that this complex must be activated, possibly through a conformation change in either one or both proteins for proper function.

Our studies further the analysis of the Wpl1p-Pds5p complex to demonstrate its regulation of cohesin function *in vivo.* This proposed role for Wpl1p in recycling cohesin, in part through its association with Pds5p, assigns a biological role for the previous finding that the Wpl1p-Pds5p complex promotes both cohesin loading and unloading *in vitro* (Murayama and Uhlmann 2015). It will be interesting to further parse how the Wpl1p-Pds5p complex regulates cohesin function differently from Wpl1p regulation independent of Pds5p. Furthermore, exploring the roles of Wpl1p and cohesin in S-phase induced DNA damage provide an exciting new direction in the cohesin field.

## Acknowledgements

We thank Rebecca Lamothe, Brett Robison and Lorenzo Costantino for critical reading of the manuscript and helpful comments. We thank the rest of the Koshland Lab and Martin Kupiec for their moral and technical support as well as fruitful discussions. We thank Kim Nasmyth for kindly sharing the *SMC3*-*MCD1* fusion construct with us. This work was supported by the NIH Grant 1R35GM118189-01 (to D.K.)

---

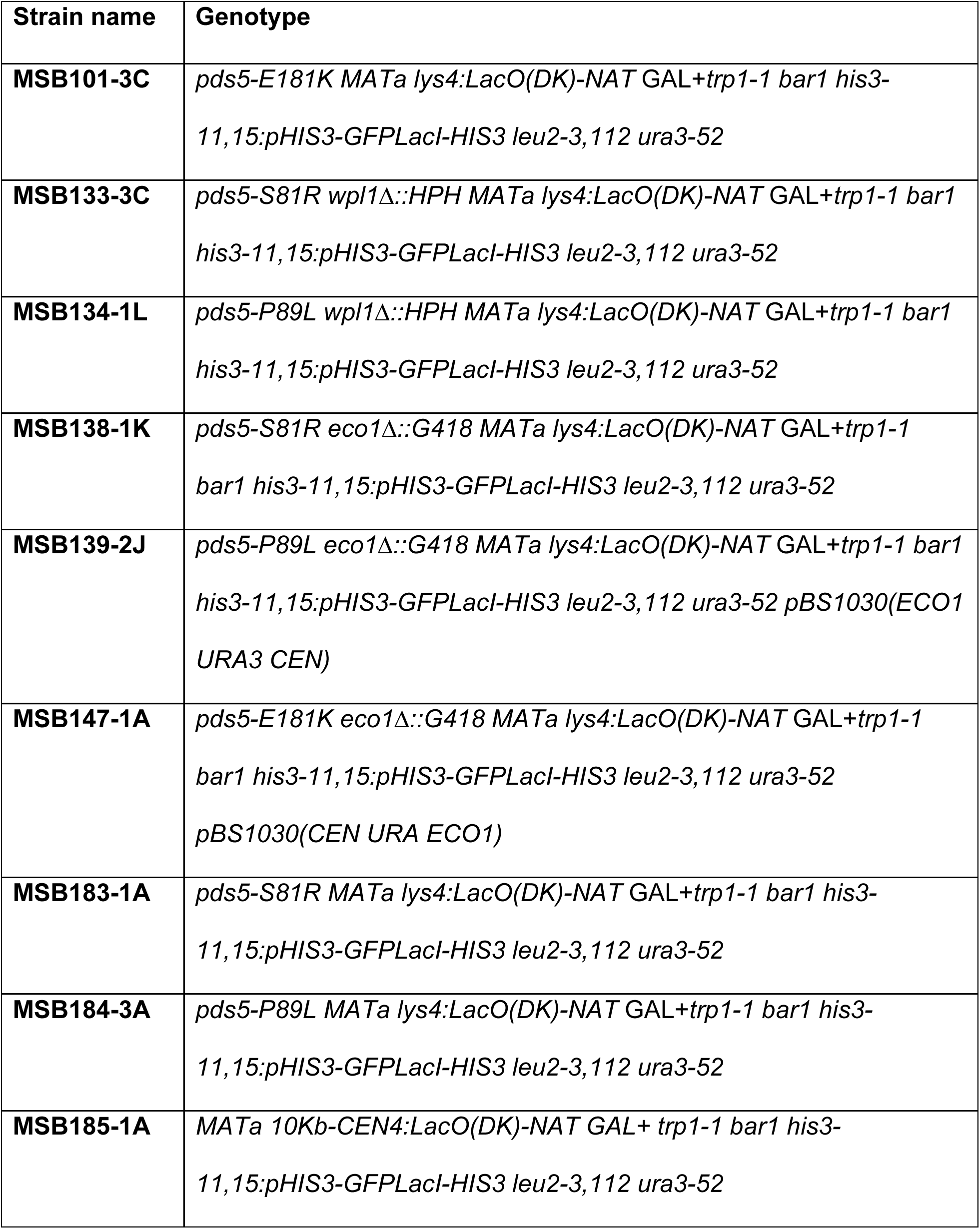

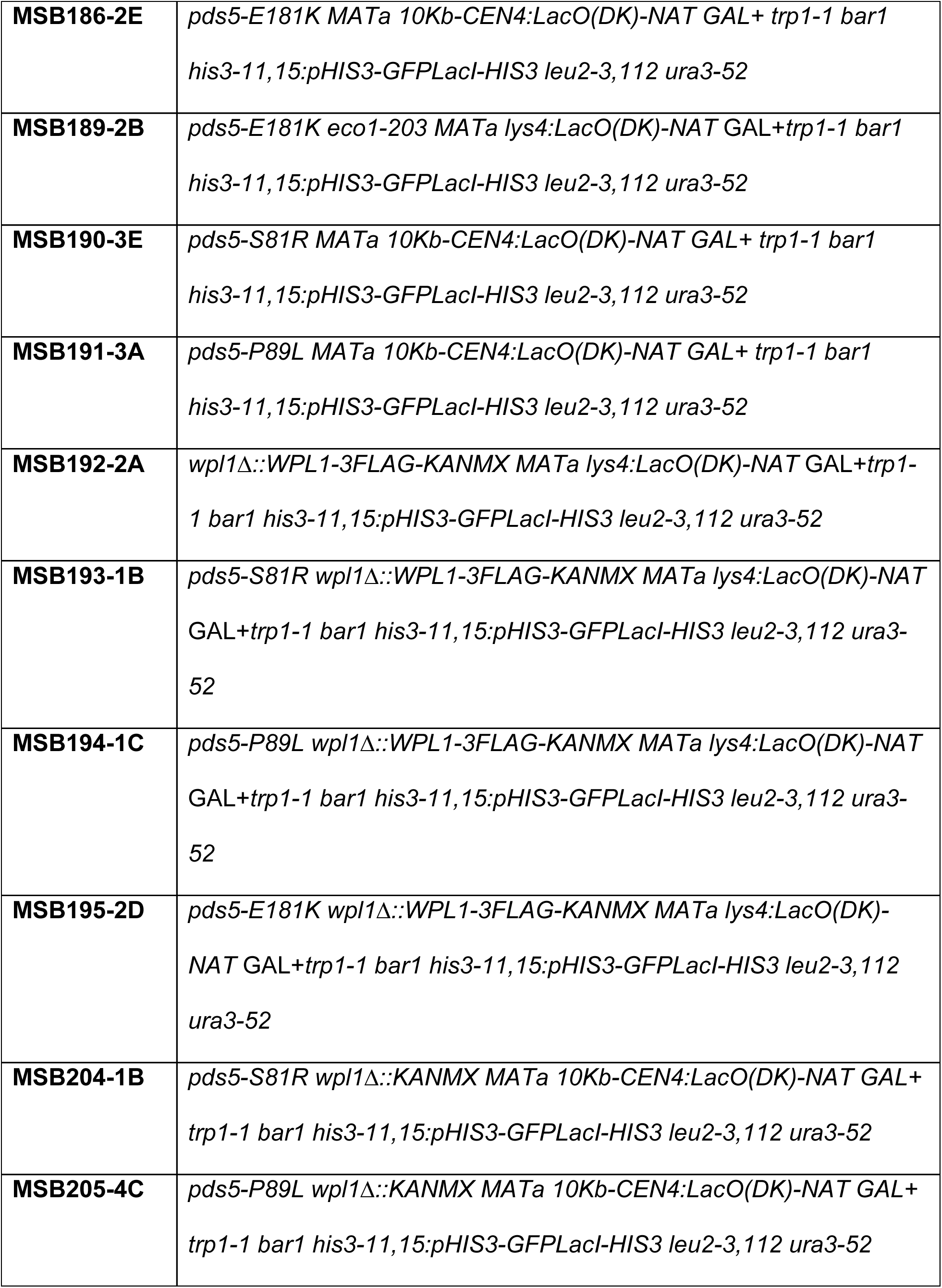

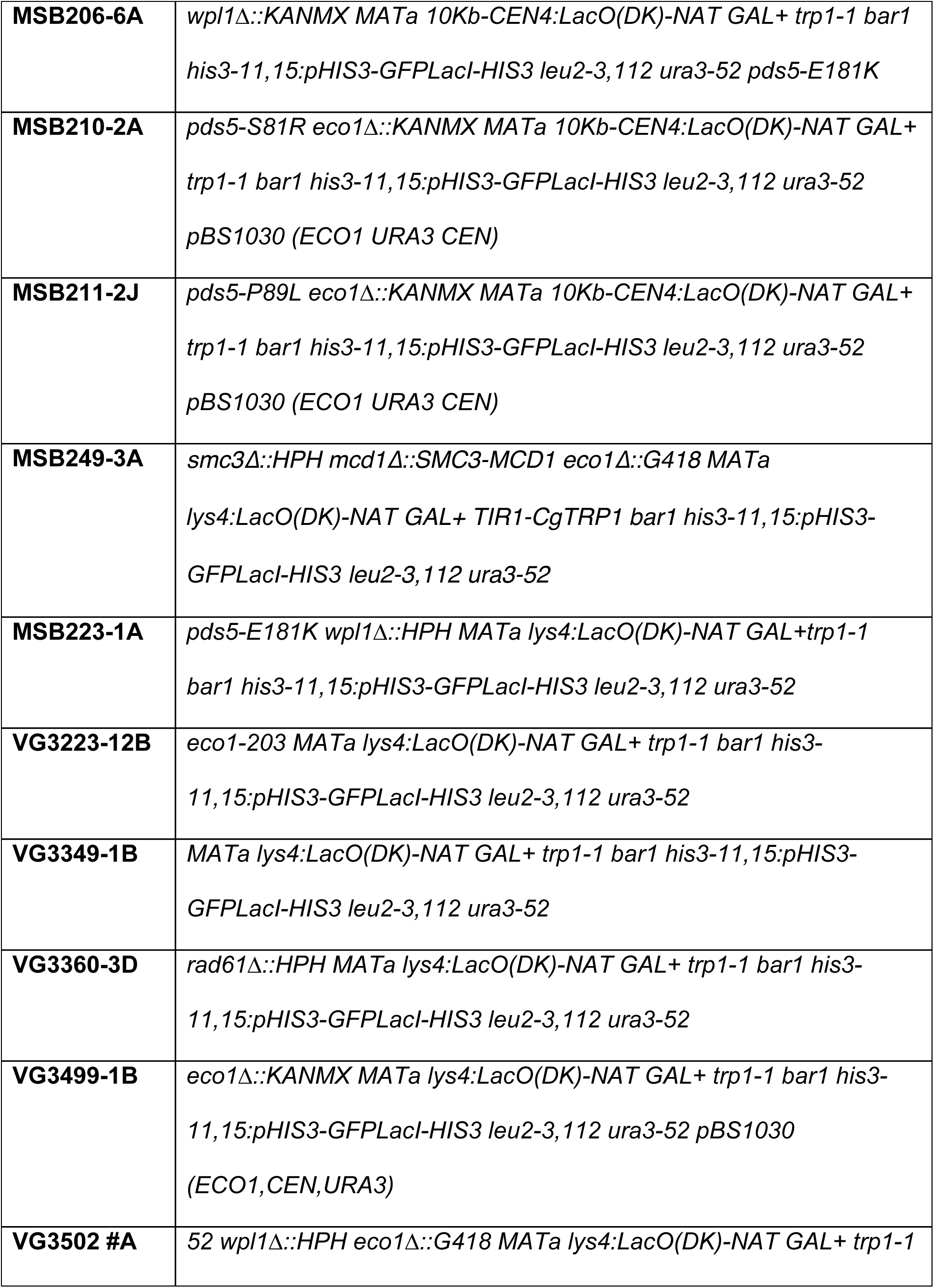

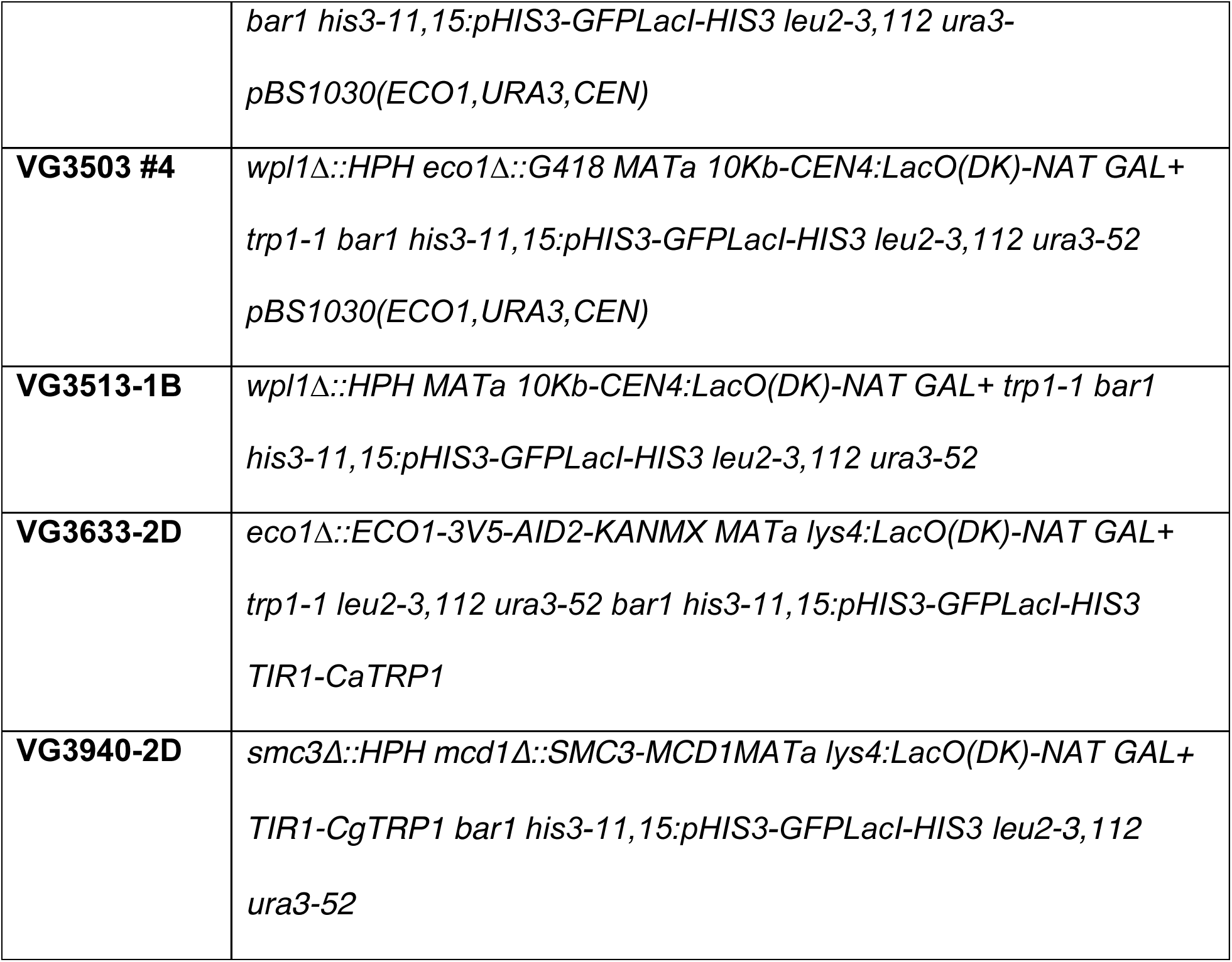
Strain list.

## References

Avemann K, Knippers R, Koller T, Sogo JM. 1988. Camptothecin, a Specific Inhibitor of Type-I Dna Topoisomerase, Induces Dna Breakage at Replication Forks. Molecular and Cellular Biology 8:3026–3034.

Beckouët F, Srinivasan M, Roig MB, Chan K-L, Scheinost JC, Batty P, Hu B, Petela N, Gligoris T, Smith AC, et al. 2016. Releasing Activity Disengages Cohesin’s Smc3/Scc1 Interface in a Process Blocked by Acetylation. Molecular Cell 61:563–574.

Buheitel J, Stemmann O. 2013. Prophase pathway-dependent removal of cohesin from human chromosomes requires opening of the Smc3-Scc1 gate. The EMBO Journal 32:666–676.

Chan K-L, Gligoris T, Upcher W, Kato Y, Shirahige K, Nasmyth K, Beckouët F. 2013. Pds5 promotes and protects cohesin acetylation. Proceedings of the National Academy of Sciences 110:13020–13025.

Chan K-L, Roig MB, Hu B, Beckouët F, Metson J, Nasmyth K. 2012. Cohesin’s DNA exit gate is distinct from its entrance gate and is regulated by acetylation. Cell 150:961–974.

Çamdere G, Guacci V, Stricklin J, Koshland D. 2015. The ATPases of cohesin interface with regulators to modulate cohesin-mediated DNA tethering. eLife 4:1–66.

Feytout A, Vaur S, Genier S, Vazquez S, Javerzat JP. 2011. Psm3 Acetylation on Conserved Lysine Residues Is Dispensable for Viability in Fission Yeast but Contributes to Eso1-Mediated Sister Chromatid Cohesion by Antagonizing Wpl1. Molecular and Cellular Biology 31:1771–1786.

Game JC, Birrell GW, Brown JA, Shibata T, Baccari C, Chu AM, Williamson MS, Brown JM. 2003. Use of a Genome-Wide Approach to Identify New Genes that Control Resistance of Saccharomyces cerevisiae to Ionizing Radiation. Radiation Research 160:14–24.

Gandhi R, Gillespie PJ, Hirano T. 2006. Human Wapl Is a Cohesin-Binding Protein that Promotes Sister-Chromatid Resolution in Mitotic Prophase. Current Biology 16:2406–2417.

Gligoris TG, Scheinost JC, Bürmann F, Petela N, Chan K-L, Uluocak P, Beckouët F, Gruber S, Nasmyth K, Lowe J. 2014. Closing the cohesin ring: structure and function of its Smc3-kleisin interface. Science 346:963–967.

Goto Y, Yamagishi Y, Shintomi-Kawamura M, Abe M, Tanno Y, Watanabe Y. 2017. Pds5 Regulates Sister-Chromatid Cohesion and Chromosome Bi-orientation through a Conserved Protein Interaction Module. Current Biology 27:1005–1012.

Gruber S, Arumugam P, Katou Y, Helmhart W, Shirahige K, Nasmyth K. 2006. Evidence that loading of cohesin onto chromosomes involves opening of its SMC hinge. Cell 127:523–537.

Gruber S, Haering CH, Nasmyth K. 2003. Chromosomal cohesin forms a ring. Cell 112:765–777.

Guacci V, Koshland D. 1994. Chromosome condensation and sister chromatid pairing in budding yeast. The Journal of Cell Biology 125:517–530.

Guacci V, Koshland D. 2012. Cohesin-independent segregation of sister chromatids in budding yeast. Molecular Biology of the Cell 23:729–739.

Guacci V, Koshland D, Strunnikov A. 1997. A direct link between sister chromatid cohesion and chromosome condensation revealed through the analysis of MCD1 in S. cerevisiae. Cell 91:47–57.

Guacci V, Stricklin J, Bloom MS, Guō, X, Bhatter M, Koshland D. (2015). A novel mechanism for the establishment of sister chromatid cohesion by the ECO1 acetyltransferase. Molecular Biology of the Cell 26:117–133.

Guacci V, Yamamoto A, Strunnikov A, Kingsbury J, Hogan E, Meluh P, Koshland D. 1993. Structure and Function of Chromosomes in Mitosis of Budding Yeast. Cold Spring Harbor Symposia on Quantitative Biology 58:677–685.

Hartman T, Stead K, Koshland D, Guacci V. 2000. Pds5p is an essential chromosomal protein required for both sister chromatid cohesion and condensation in Saccharomyces cerevisiae. The Journal of Cell Biology 151:613–626.

Hartwell LH. 1974. Saccharomyces cerevisiae cell cycle. Bacteriological Review 38:164–198.

Heidinger-Pauli JM, Mert O, Davenport C, Guacci V, Koshland D. 2010. Systematic reduction of cohesin differentially affects chromosome segregation, condensation, and DNA repair. Current Biology 20:957–963.

Kueng S, Hegemann B, Peters BH, Lipp JJ, Schleiffer A, Mechtler K, Peters J-M. 2006. Wapl Controls the Dynamic Association of Cohesin with Chromatin. Cell 127:955–967.

Lopez-Serra L, Lengronne A, Borges V, Kelly G, Uhlmann F. 2013. Budding yeast Wapl controls sister chromatid cohesion maintenance and chromosome condensation. Current Biology 23:64–69.

Murayama Y, Uhlmann F. 2015. DNA Entry into and Exit out of the Cohesin Ring by an Interlocking Gate Mechanism. Cell 163:1628–1640.

Noble D, Kenna M, Dix M, Skibbens RV, Unal E. 2006. Intersection between the regulators of sister chromatid cohesion establishment and maintenance in budding yeast indicates a multi-step mechanism. Cell Cycle 5:2528–2536.

Onn I, Heidinger-Pauli JM, Guacci V, Unal E, Koshland D. 2008. Sister Chromatid Cohesion: A Simple Concept with a Complex Reality. Annual Review of Cell and Developmental Biology 24:105–129.

Ouyang Z, Zheng G, Tomchick DR, Luo X, Yu H. 2016. Structural Basis and IP6 Requirement for Pds5-Dependent Cohesin Dynamics. Molecular Cell 62:248–259.

Rolef Ben-Shahar T, Heeger S, Lehane C, East P, Flynn H, Skehel M, Uhlmann F. 2008. Eco1-dependent cohesin acetylation during establishment of sister chromatid cohesion. Science 321:563–566.

Rowland BD, Roig MB, Nishino T, Kurze A, Uluocak P, Mishra A, Beckouët F, Underwood P, Metson J, Imre R, et al. 2009. Building Sister Chromatid Cohesion: Smc3 Acetylation Counteracts an Antiestablishment Activity. Molecular Cell 33:763–774.

Saleh-Gohari N, Bryant HE, Schultz N, Parker KM, Cassel TN, Helleday T. 2005. Spontaneous Homologous Recombination Is Induced by Collapsed Replication Forks That Are Caused by Endogenous DNA Single-Strand Breaks. Molecular and Cellular Biology 25:7158–7169.

Shintomi K, Hirano T. 2009. Releasing cohesin from chromosome arms in early mitosis: opposing actions of Wapl-Pds5 and Sgo1. Genes & Development 23:2224–2236.

Skibbens RV, Corson LB, Koshland D, Hieter P. 1999. Ctf7p is essential for sister chromatid cohesion and links mitotic chromosome structure to the DNA replication machinery. Genes & Development 13:307–319.

Stead K, Aguilar C, Hartman T, Drexel M, Meluh P, Guacci V. 2003. Pds5p regulates the maintenance of sister chromatid cohesion and is sumoylated to promote the dissolution of cohesion. The Journal of Cell Biology 163:729–741.

Ström L, Lindroos HB, Shirahige K, Sjögren C. 2004. Postreplicative Recruitment of Cohesin to Double-Strand Breaks is Required for DNA Repair. Molecular Cell 16:1003–1015.

Strumberg D, Pilon AA, Smith M, Hickey R, Malkas L, Pommier Y. 2000. Conversion of topoisomerase I cleavage complexes on the leading strand of ribosomal DNA into 5′-phosphorylated DNA double-strand breaks by replication runoff. Molecular and Cellular Biology 20:3977–3987.

Sutani T, Kawaguchi T, Kanno R, Itoh T, Shirahige K. 2009. Budding Yeast Wpl1(Rad61)-Pds5 Complex Counteracts Sister Chromatid Cohesion-Establishing Reaction. Current Biology 19:492–497.

Tanaka K, Hao Z, Kai M, Okayama H. 2001. Establishment and maintenance of sister chromatid cohesion in fission yeast by a unique mechanism. The EMBO Journal 20:5779–5790.

Tittel-Elmer M, Lengronne A, Davidson MB, Bacal J, François P, Hohl M, Petrini JHJ, Pasero P, Cobb JA. 2012. Cohesin Association to Replication Sites Depends on Rad50 and Promotes Fork Restart. Molecular Cell 48:98–108.

Tong K, Skibbens RV. 2014. Cohesin without Cohesion: A Novel Role for Pds5 in Saccharomyces cerevisiae. PLoS ONE 9:e100470–14.

Unal E, Arbel-Eden A, Sattler U, Shroff R, Lichten M, Haber JE, Koshland D. 2004. DNA damage response pathway uses histone modification to assemble a double-strand break-specific cohesin domain. Molecular Cell 16:991–1002.

Unal E, Heidinger-Pauli JM, Kim W, Guacci V, Onn I, Gygi SP, Koshland D. 2008. A Molecular Determinant for the Establishment of Sister Chromatid Cohesion. Science 321:566–569.

Zhang J, Shi X, Li Y, Kim B-J, Jia J, Huang Z, Yang T, Fu X, Jung SY, Wang Y, et al. 2008. Acetylation of Smc3 by Eco1 is required for S phase sister chromatid cohesion in both human and yeast. Molecular Cell 31:143–151.

Zhou L, Liang C, Chen Q, Zhang Z, Zhang B, Yan H. 2017. The N-Terminal NonKinase-Domain-Mediated Binding of Haspin to Pds5B Protects Centromeric Cohesion in Mitosis. Current Biology 27:992–1004.

